# scFPC-DE: Robust Differential Expression Analysis Along Single Cell Trajectories via Functional Principal Component Analysis

**DOI:** 10.1101/2025.11.03.686374

**Authors:** Ricardo J. López Candelaria, Yu Qian, Fang Chen, Mansun Law, Yun Zhang, Xing Qiu

## Abstract

**Motivation:** Identifying temporally differentially expressed genes (TDEGs) along pseudotime trajectories from single cell RNA sequencing (scRNA-seq) data helps characterize the cellular states that underlie the dynamic process of cellular development. However, existing tests based on generalized additive models (GAMs) suffer from increased false positive rates under zero inflation caused by high dropouts, a ubiquitous technical artifact of scRNA-seq data. Furthermore, by testing each gene independently, existing tests ignore the variance-covariance structure shared across genes along the trajectory, leading to suboptimal power and reduced interpretability.

**Results:** We present scFPC-DE, a trajectory-based differential expression analysis (TDEA) method based on functional data analysis (FDA). It models the gene expression as a function of pseudotime in the *L*^2^ space and represents the covariance structure of these functions by eigenfunctions derived from functional principal component (FPC) analysis. This approach effectively captures informative gene expression patterns along the trajectory, while mitigating the influence of zero inflation in both simulation and real data analysis. In simulations, scFPC-DE exhibited superior control of type I error and achieved the highest ROC-AUC among competing methods. When applied to an scRNA-seq dataset of B cell subtypes, scFPC-DE uniquely identified TDEGs enriched for B cell differentiation pathways, outperforming existing methods in biological relevance. These results show that scFPC-DE effectively captures the shared gene expression variation and pseudo-temporal structure along the single cell trajectory for TDEG identification.

**Availability:** R package and code vignettes are publicly available at https://github.com/LopezRicardo1/scFPCDE.

**Supplementary information:** Supplementary data are available at *Bioinformatics* online

## Introduction

Single cell RNA sequencing (scRNA-seq) has revolutionized transcriptomics by enabling the measurement of gene expression at the resolution of individual cells, capturing cellular heterogeneity and revealing complex biological processes such as cell differentiation [1]. scRNA-seq experiments generate large-scale datasets composed of thousands of cells and genes, posing substantial statistical and computational challenges for data processing and analysis. Gene expression signals in single cells are much lower than those in bulk transcriptomics. Robust statistical methods are required to effectively capture the characteristics of cells and extract meaningful biological insights from scRNA-seq data. An emerging, rapidly growing area of scRNA-seq analysis is single cell trajectory inference, which aims to reconstruct the continuous development of cells through dynamic biological processes such as cell differentiation and cell activation [1]. This is typically achieved by estimating a latent variable known as pseudotime using methods such as *Monocle3* [2], which orders cells along a trajectory that reflects the biological states of the cells in the process. Trajectory-based differential expression (DE) analysis aims to identify temporally differentially expressed genes (TDEGs) along a cell trajectory inferred via pseudotime. Methods for trajectory-based DE analysis, such as the “regression analysis” approach in the DE module in *Monocle3* [3] and *tradeSeq* [4], use generalized additive models (GAMs) of smooth functions for fitting gene expression values along the estimated pseudotime. There is also a “graph-autocorrelation analysis” approach for DE analysis in *Monocle3* that uses Moran’s I [5] to detect genes with expression patterns spatially correlated with the reconstructed cell graph/trajectory. While this test is sensitive to localized or non-monotonic patterns, it does not provide effect estimates or easy covariate adjustment (including experimental designs). In addition, its graph reconstruction, gene-wise Moran’s I calculations, and Monte-Carlo resampling for *p*-value calculation scale very poorly with cell and gene counts. PseudotimeDE [6] accounts for uncertainties in pseudotime estimation and applies a permutation-based inference framework with the GAM (Gaussian or negative-binomial) approach to produce recalibrated *p*-values. Because it corrects for pseudotime variability and has outperformed alternatives such as *Monocle3*-DE and *tradeSeq* in recent benchmarks, PseudotimeDE is widely regarded as the current state-of-the-art for TDEG detection.

Most of these modeling approaches rely on parametric assumptions, such as negative binomial or zero-inflated negative binomial distributions for the scRNA-seq count data. However, parametric approaches are vulnerable to deviations from the underlying distribution assumptions. An alternative is to model the gene expression changes along pseudotime using functional principal component analysis (FPCA), a well-known technique in functional data analysis (FDA), [7, 8]. An FPCA-based F-statistic was introduced for DE testing in time-course bulk RNA-seq studies [9], which outperformed alternative time-course methods, particularly when the number of time points is sparse or irregular. FPCA modeling was also used for robust gene set enrichment analysis (GSEA) in time-course bulk RNA-seq studies [10]. A major strength of FPCA lies in its ability to learn a data-driven basis in the *L*^2^ space (eigenfunctions of the covariance operator), which provides a representation of the dominant temporal patterns of gene expressions along time or pseudotime. This stands in contrast to approaches like GAM, which typically rely on fixed, predefined basis functions. By the celebrated Karhunen–Loève theorem [11–13], top *L* eigenfunctions estimated by FPCA capture more of the total *L*^2^ variance than any other *L*-dimensional basis, making them the most efficient parsimonious low-rank representation of temporal data. In the scRNA-seq context, where each cell provides a pseudo-temporal observation, FPCA naturally captures the *<u>shared temporal patterns</u>* across genes and pseudotime points (i.e., cells along the trajectory), overcoming the time point sparsity in the bulk RNA-seq experiments that often involve a limited number of discrete sampling times.

Unlike bulk RNA-seq, scRNA-seq data are known for high zero inflation. Zero counts in scRNA-seq data may arise from sampling zero, technical zero, and biological zero [14]. The dropout technical artifact due to the low amount of mRNA in scRNA-seq experiments contributes to excessive zero counts in scRNA-seq data [15]. High zero inflation presents a non-trivial challenge for downstream analysis because many standard approaches assume continuous, well-sampled observations. Consequently, single cell trajectory analysis requires methods that remain robust when a large proportion of measurements are zeros.

Motivated by these considerations, we introduce scFPC-DE, a trajectory-based DE analysis (TDEA) testing method grounded in the FDA hypothesis testing principles [16]. The traditional functional F-test (FPC-F) summarizes trajectory variability by the ratio of between- to within-trajectory variance. When zeros dominate a gene’s profile, the denominator can be very small, producing spuriously large F-statistic value driven by just a few nonzero points. In contrast, scFPC-DE avoids this issue by using a distance-based test statistic (*L*^2^ distance in the subspace spanned by eigenfunctions) which is numerically stable even when expression profiles contain excessive zeros. Secondly, scFPC-DE constructs its functional basis (eigenfunctions) from a selected subset of trajectory-informative genes (TIGs): Well-expressed genes that exhibit sufficient variation relevant to the pseudotime. As a result, our trajectory model reflects biologically meaningful variation, leading to more robust inference and substantially tighter false-positive control in zero-inflated scRNA-seq data. Our strategy addresses the limitations of existing methods and provides a powerful framework for cell trajectory downstream analysis.

In our simulation studies designed to reflect the high zero inflation nature of scRNA-seq data, scFPC-DE stringently controls the type I error at the nominal level, while competing methods exhibited considerable inflation of type I error. scFPC-DE also achieved comparable or higher power than existing approaches. Consequently, scFPC-DE attained the highest area under the curve (AUC) in Receiver Operating Characteristic (ROC) analysis among the methods tested.

We further evaluate method performance using a real scRNA-seq dataset of phenotypically sorted B cell subtypes [17]. scFPC-DE and PseudotimeDE were applied to identify TDEGs associated with the B cell differentiation trajectory of transitional, naïve, and classical memory B cell subtypes. We apply *ReactomePA* (for pathway enrichment analyses) [18], and the *Enrichr-KG* online tool (for phenotype/network queries) [19], to evaluate biological relevance of the TDEGs identified by both methods. Compared with PseudotimeDE, the TDEGs identified by scFPC-DE form a more concise list that exhibits greater biological relevance and stronger enrichment for pathways related to B cell development and immune system processes. This more stringent and focused DEG detection using scFPC-DE is consistent with the tight type I error control observed in simulations, which supports more targeted and biologically interpretable downstream analyses.

The remainder of this manuscript is organized as follows. Section 2 develops the core statistical properties of the FPC model for trajectory gene expression and introduces the distance-based test statistic in the FPC latent space. Section 3 presents results from zero-inflated simulation studies and compares the performance of scFPC-DE with previously established FPC-F and PseudotimeDE methods, using the latter as the current benchmark for trajectory-based differential expression analysis (TDEA). This section also includes results from the real B cell scRNA-seq dataset and enrichment analysis of the identified TDEGs. Section 4 concludes with a discussion of the contributions. The scFPC-DE R package is publicly available at GitHub: https://github.com/LopezRicardo1/scFPCDE.

## Methods

Functional principal component analysis (FPCA) [7, 20] is an extension of standard PCA for continuous functions. Here, we propose to model the gene expression changes along the trajectory as a function of pseudotime and use FPCA to capture the main sources of variations across the gene expression functions, providing a data-driven framework for modeling the gene expression changes along the pseudotime trajectory.

### 2.1 Modeling Cell Trajectories with FPCA

Let *Y_i_*(*t_j_*) denote the log-transformed, centered gene expression of gene *i* at pseudotime point *j*. Using FPCA, we can express them as

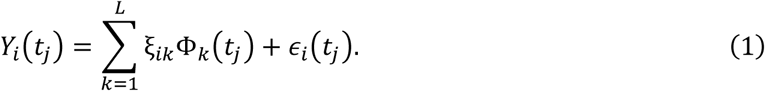

Here Φ*_k_*(*t*), *k* = 1, … *L*, are the top *L* eigenfunctions that capture the dominant temporal expression modes, and *ξ_ik_* are the corresponding functional principal component (FPC) scores. The residual term *∈_i_*(*t_j_*) models random noise and accounts for the remaining variability. FPC scores are obtained by projecting each gene trajectory onto the linear space spanned by eigenfunctions learned non-parametrically satisfying

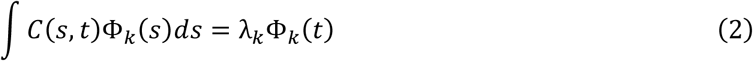

where *C*(*s*, *t*) and *λ_k_* denotes the pseudotime covariance and *k*-th eigenvalue, respectively (see Supplementary Section 1.1). Genes with large FPC scores have larger pseudo-temporal variations that align with the dynamic behavior represented by eigenfunctions, while small scores indicate flat or noise-driven profiles. We performed FPCA using a standard procedure based on roughness penalized B-spline [7, 21], implemented in the *fda* R package [21]

### 2.2 Trajectory-based Differential Expression FPC Test

Under the null hypothesis of no pseudo-temporal variation, FPC scores remain centered at zero with their variability driven only by the measurement noise, yielding small magnitudes of FPC scores relative to the alternative hypothesis. Formally, for each gene *i*, the null and alternative hypotheses can be stated as:

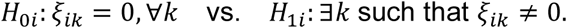

We use the squared Euclidean norm of FPC scores as the test statistics:

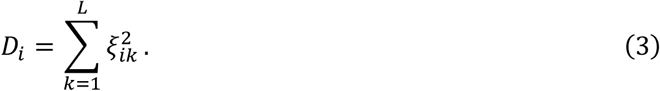

Mathematically, *D_i_* is the squared *L*^2^-distance between the constant zero function and the *i*th trajectory projected to the linear subspace spanned by the eigenfunctions. A larger *D_i_* indicates stronger evidence of the pseudotime dynamics. To obtain a null distribution of *D_i_*, we randomly permute the pseudotime labels across cells and recompute *D_i_* from these permutations. The permutation-based empirical p-value is defined as:

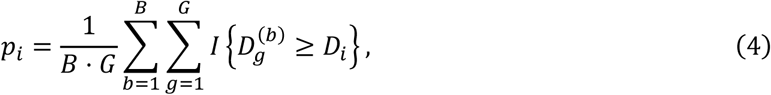

where 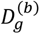 is the test statistic for gene *g* under the *b*-th permutation, and *B*, *G* are the number of permutations and genes, respectively. A significant gene identified by this procedure is called a temporally differentially expressed gene (TDEG).

An outline of the scFPC-DE workflow is provided in Figure 1. We first apply an established trajectory analysis tool (e.g., *Monocle3*) to cluster cells and estimate pseudotime from properly processed scRNA-seq data. Next, we model each gene’s log-transformed, mean-centered expression as a smooth function of pseudotime using roughness-penalized B-splines and perform FPCA to extract top eigenfunctions and corresponding FPC scores. For each gene, we compute the distance statistic *D_i_* with and without random pseudotime permutations. The resulting null distribution of *D_i_* enables us to derive empirical *p*-values and set a decision boundary in the FPC score space to identify TDEGs. Smoothed gene expression trajectories produced by scFPC-DE can be visualized in the FPC space overlaid with a directional field from their pseudo-temporal derivatives. Finally, we perform gene-set and pathway enrichment analyses on the identified TDEGs to understand the underlying biological processes.

**Fig. 1.**
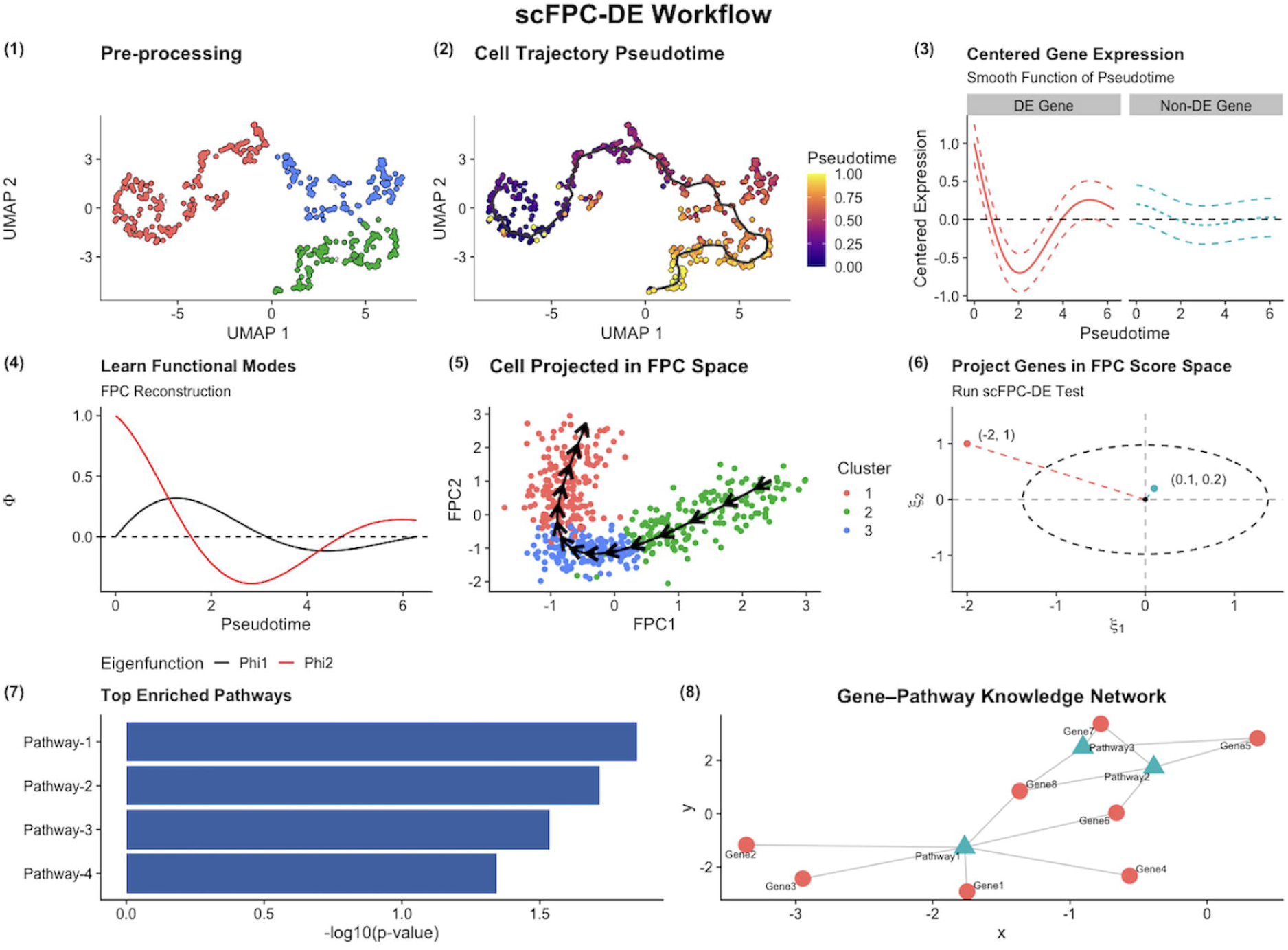
scFPC-DE workflow. (1) Data preprocessing of the input scRNA-seq dataset, including normalization, log-transformation, dimensionality reduction, finding neighbors, UMAP/tSNE embedding, and clustering. Note that the scFPC-DE workflow uses the log-transformed gene expression data as the input data for the downstream analysis workflow. (2) Trajectory analysis for estimating the pseudotime of cells in a trajectory of interest. (3) Modeling the log-transformed gene expression as a smoothed function of pseudotime centered at functional mean expression of 0 for each gene. Examples of a temporally DE gene (left) and a temporally non-DE gene (right). (4) Identifying the top eigenfunctions *Φ_k_*(*t*) from FPCA and construction of the FPC latent space. (5) Visualization of the reconstructed cell trajectory in the FPC space. (6) Projecting each gene onto the FPC space allows calculation the distance statistic *D_i_* for hypothesis testing. Genes are represented as coordinates with respect to the eigenfunctions, with the significance decision boundary is denoted by dashed ellipsoid; genes positioned farther away are classified as TDEGs/TIGs. (7-8) Enrichment analysis of DEGs and knowledge network visualization using online tools.

### 2.3 FPC Refinement with Trajectory-Informative Genes

Data from scRNA-seq experiments are highly zero-inflated due to the technical artifact of dropouts [14, 15]. To mitigate the excessive sparsity in the data that may dilute the estimation of pseudo-temporal expression patterns, we adopt a strategy to estimate the eigenfunctions by focusing on a subset of trajectory-informative genes (TIGs) that are likely to exhibit non-flat, informative pseudo-temporal trends.

Selecting highly variable genes is a common practice in the clustering-based scRNA-seq data analysis pipeline (e.g., *Seurat* [22]). However, standard approaches based on dispersion metrics do not necessarily capture useful variability along the pseudotime trajectory. To ensure our FPCA basis represents genuinely dynamic genes, we implement a two-stage procedure focusing on TIGs. First, we fit an initial FPCA model to all genes and compute the distance statistic *D_i_* for each gene, which captures its overall pseudo-temporal variation. Empirical evidence suggests the eigenfunctions obtained in this step are overly smoothed, likely due to the “dilution effects” of non-informative genes and zero inflation (see Supplementary Section 3.2; Figure 8). Next, we select top 20~30% genes ranked by *D_i_* as TIGs and re-estimate the covariance surface, eigenfunctions, and FPC scores using only TIGs. The result is a stable functional basis that reflects major temporal dynamics of informative genes. This retraining procedure improves interpretability and the statistical power of our DE test, particularly under high sparsity.

### 2.4 Simulation Study

We use *scDesign3* [23] to generate synthetic data that reflect realistic characteristics of scRNA-seq data. Simulated dataset is designed to consist of 4,000 genes, including 500 true TDEGs along 500 pseudotime points (cells) equally spaced in a unit interval.

To model the dynamic expression patterns, we design the simulation model for the centered and log-transformed mean gene expression curves using three basis functions with distinct dynamic patterns (see Supplementary Figure 1):

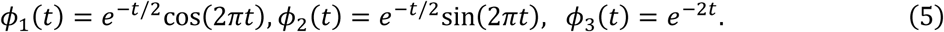

FPC scores for each TDEG are sampled from a standard normal distribution 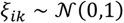, scaled by eigenvalues prioritizing dominant expression trends: *λ*_1_ = 1, *λ*_2_ = 0.5, and *λ*_3_ = 0.25. For each TDEG, the true mean expression at each pseudotime point is defined as:

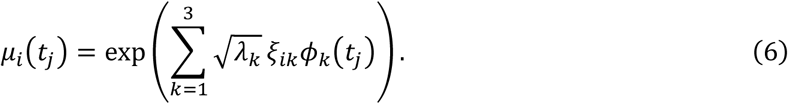

Non-TDEGs have constant mean values along pseudotime, sampled uniformly between 0 and the median expression of all simulated TDEGs.

Gene expression counts are first generated using a programmatic random number generation from a zero-inflated negative binomial (ZINB) distribution defined as 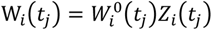, where 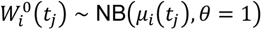, and *Z_i_*(*t_j_*) ~ Bernoulli(*p* = 0.25) as a dropout indicator introducing high zero inflation at the same level as observed in the real data analysis. Using the programmatically generated count data as input, we then use *scDesign3* to further introduce stochastic components that reflect the realistic characteristics of real scRNA-seq data. The *scDesign3* simulated data no longer follows the classical distributional assumptions, thus limiting the validity of methods based on the GAM approach. The high zero inflation dataset will serve as the primary simulation dataset in this manuscript. In addition, we generated three supplementary simulation datasets with low to moderate zero inflation in Supplementary Section 2.1, and present the results in Supplementary Section 2.2, which show the progressive impact of zero inflation on the method performance.

### 2.5 Real Data Analysis

We applied scFPC-DE to a publicly available real scRNA-seq dataset of B cell subtypes [17] downloaded from the EMLB-EBI ArrayExpress data portal (https://www.ebi.ac.uk/biostudies/arrayexpress) with Array Express accession: E-MTAB-9544. We preprocessed the raw counts with the Seurat workflow [24] and then inferred cell trajectories and pseudotime ordering with *Monocle3* [3]. The resulting pseudotime trajectories are consistent with those reported in [17]. For trajectory-based differential expression analyses, we focused on the developmental trajectory of transitional (Trans), naïve (Naïve), and classical memory (C-mem1) B cell clusters. Genes with exceptionally low expression or high sparsity across these subsets were removed (see Supplementary Figure 7). The filtered data still comprises 93.9% overall sparsity (Supplementary Table 3), mirroring the sparsity of our primary simulated data.

Using scFPC-DE, we identified consistent TDEGs including the canonical markers found in the original study [17] and B-cell specific pathways. All analyses can be reproduced using scripts available at: https://github.com/LopezRicardo1/scFPCDE/tree/main/vignettes.

We performed gene set enrichment analysis using the *ReactomePA* [18] package and Enrichr Knowledge Graph online platform for querying phenotype/network features from multiple libraries (https://maayanlab.cloud/enrichr-kg) [19]. Reactome pathway analysis was conducted via Over-Representation Analysis (ORA), using the sets of TDEGs identified by scFPC-DE and PseudotimeDE as input, against a common background gene universe [18]. *ReactomePA* was used for Reactome pathway enrichment, while Enrichr-KG was employed to analyze Mammalian Phenotype Level 4 ontology (2021 release) and construct phenotype-level interaction networks. Pathways and phenotype categories with FDR-adjusted p-values below 0.05 were considered significantly enriched.

## Results

### 3.1 Benchmarking Performance of scFPC-DE in Simulations

PseudotimeDE using a negative binomial generalized additive model (NB-GAM) is considered the state-of-the-art method for TDEG identification. To benchmark the method performance in simulations, we compared scFPC-DE with PseudotimeDE using NB-GAM. We also compared scFPC-DE with FPC-F, which serves as a baseline for a generic FPC-based functional test. Additional comparisons with the Gaussian and ZINB-GAM variants of PseudotimeDE are provided in Supplementary Section 2.2.

In the simulation benchmarking, we used the oracle pseudotime simulated in the synthetic data, which provides a fair performance evaluation of the TDEG identification itself and removes the uncertainty from pseudotime estimation. Table 1 summarizes the type I error rate and statistical power of the three benchmarked methods using the high zero inflation simulated dataset. scFPC-DE consistently showed the best type I error control while maintaining competitive power compared to the other two methods. Figure 2 presents the systematic evaluation of the simulation results in the ROC analysis. For type I error control, scFPC-DE always had the closest observed false positive rate as the expected nominal α level (Figure 2a). The ROC analysis also confirmed that scFPC-DE outperformed the competing methods by achieving the highest AUC statistic (Figure 2b), indicating the overall improved performance of sensitivity and specificity. For the supplementary simulated data with moderate-to-low zero inflation, the performance gap between scFPC-DE and benchmarked methods are smaller (Supplementary Section 2.2), suggesting that the differences in type I error control are tightly linked to the dropout rate in the data [14].

**Fig. 2.**
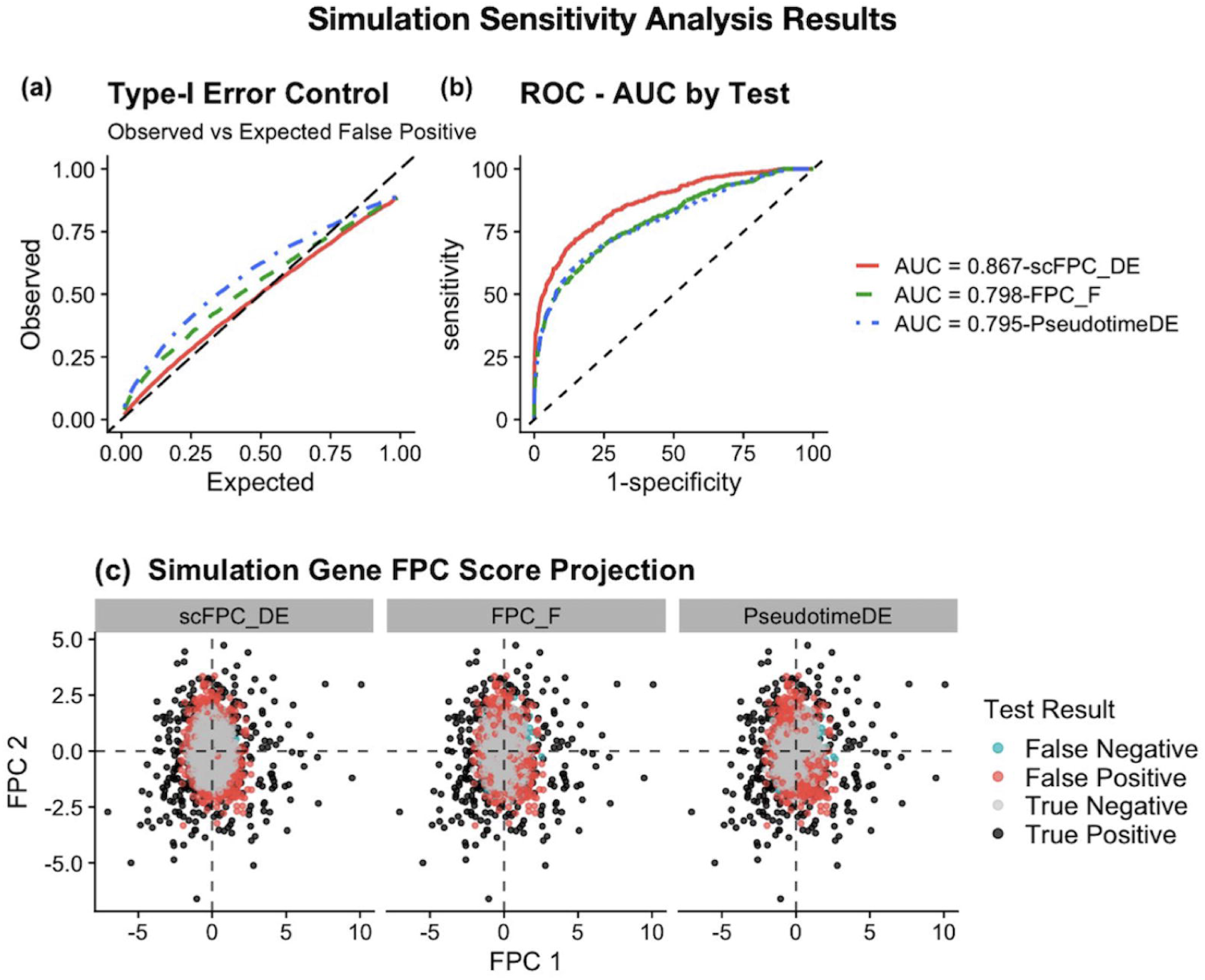
Simulation results for scFPC-DE and benchmarking methods. (a) Observed versus expected type I error curves demonstrate that scFPC-DE aligns closely with the diagonal line, demonstrating its accurate type I error control across significance thresholds. In contrast, both FPC-F and PseudotimeDE deviate above the diagonal line, indicating inflated error rates, with PseudotimeDE showing the most severe inflation. (b) ROC curve comparison among scFPC-DE (AUC = 0.867), FPC-F (AUC = 0.798), and PseudotimeDE (AUC = 0.795) shows that scFPC-DE achieves the best performance in sensitivity and specificity. (c) In the FPC projected score space, null genes (grey) cluster near the origin, while true DEGs (black) separate outward. Controlling at the 5% significance level, scFPC-DE produces fewer false positives (245 out of 4000; red), mostly located at the boundary between null and alternative regions, compared to the substantially higher false positive misclassifications observed in both FPC-F (430) and PseudotimeDE (501).

**Table 1.** Type I error rate (TIER) and statistical power (PWR) across varying significance levels (α) under a high zero inflation setting (93.9% sparsity). Simulation benchmarking results compare the performance of scFPC-DE, FPC-F, and PseudotimeDE for identifying the true TDEGs. scFPC-DE consistently exhibits the lowest TIER across all α thresholds, indicating superior type I error control. Notably, at α = 0.01, scFPC-DE reduces the TIER to 0.017, outperforming both FPC-F (0.039) and PseudotimeDE (0.049). At more lenient thresholds (α = 0.05 and 0.10), the FPRs of competing methods increase substantially, while scFPC-DE maintains markedly better calibration. Despite this advantage in error control, scFPC-DE does not compromise on statistical power. It attains power values that are comparable or superior across all tested levels—demonstrating robust sensitivity to true signal even under extreme dropout conditions. The best results for each significance level are shown in boldface.

| $\alpha$ | scFPC-DE | | FPC-F | | PseudotimeDE | |
| --- | --- | --- | --- | --- | --- | --- |
|  | TIER | PWR | TIER | PWR | TIER | PWR |
| 0.01 | **0.017** | **0.434** | 0.039 | 0.406 | 0.049 | 0.426 |
| 0.05 | **0.070** | 0.602 | 0.123 | 0.552 | 0.143 | **0.604** |
| 0.1 | **0.130** | **0.694** | 0.193 | 0.632 | 0.217 | 0.674 |

To further illustrate from where scFPC-DE gained the overall improvement, we projected the simulated data in the FPC score space (Figure 2c). By employing FPCA to capture the global covariance patterns across genes [9], scFPC-DE is able to define a more precise and stable null boundary in the FPC score space than the gene-wise GAM approaches [4, 25]. That is the main reason scFPC-DE achieves superior discrimination between true signal and noise (Figure 2c).

In the simulation studies, we also compared the computing time needed by each method. scFPC-DE demonstrated substantial computational gains, requiring only 4.82 seconds on average per dataset—more than ten times faster than the computing time of PseudotimeDE (49.62 seconds; Supplementary Table 2).

### 3.2 scFPC-DE Performance in Real Data Analysis

In the scRNA-seq dataset of B cell subtypes [17], we reconstructed the pseudotime trajectory using *Monocle3* (Figure 3a–b), which reproduces the results of Figure 2A in the original study. We captured the cell trajectory of the development pathway of Trans, Naïve, and C-mem1 B cell clusters, which recapitulates the biological cell states along the pseudotime (Figure 3c). By applying scFPC-DE, gene expression changes along the trajectory are modeled as smooth functions of pseudotime. In Figure 3d, we visualized selected genes for the Trans, Naïve, and C-mem 1 trajectory from the original study using our scFPC-DE framework. As expected, all selected genes were identified as TDEG using scFPC-DE.

**Fig. 3.**
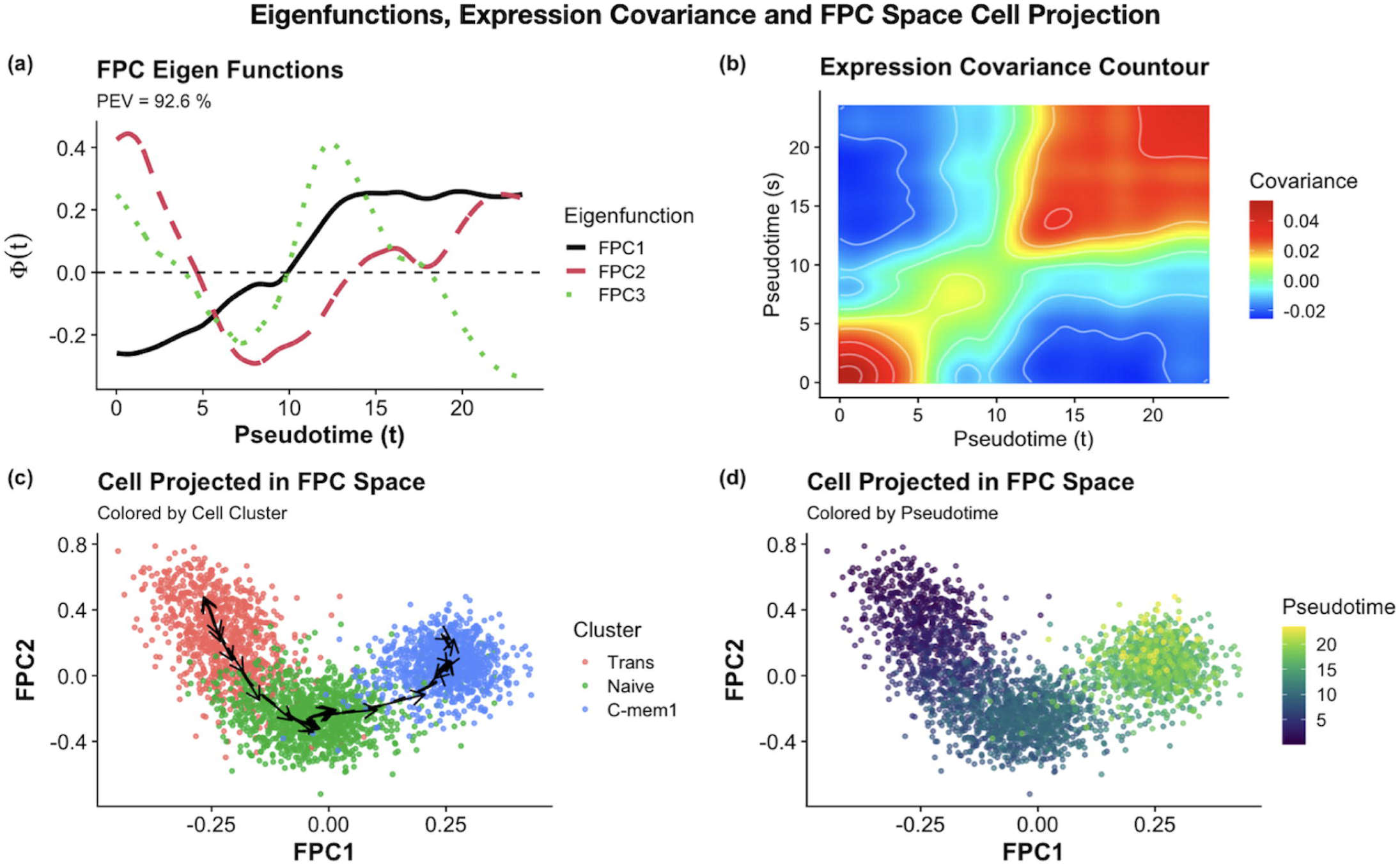
Pseudotime reconstruction and smoothed gene expression functions using real data. (a) 3-D UMAP (UMAP 1 and UMAP 3) as referenced in Stewart et al. (2021), with reconstructed *Monocle3* cell trajectory. (b) Pseudotime-colored UMAP shows consistency with the results shown in referenced study. (c) Cell type ordering and pseudotime distribution for the Trans, Naive, and CMem1 subtypes annotated by the referenced study. (d) Gene expression functions for the biologically relevant genes identified in Stewart et al. (2021), Figure 2C. The smoothed gene expression functions were modeled by the FPC framework. scFPC-DE also identified these genes as TDEGs after false discovery rate (FDR) correction.

In the FDA-based FPC modeling framework, FPCA-derived eigenfunctions (i.e., FPCs) can effectively capture the principal modes of gene expression variations along the pseudotime (Figure 4a). The functional variance-covariance structure, *C*(*t*, *s*), shared among genes indicate that there are increased gene-gene interactions among neighboring pseudotime points (Figure 4b). The linear structure of eigenfunctions also permits efficient computation of time derivatives, which may suggest the directions of cell development in the directional vectors in the FPC score space (Figure 4c). These vectors, derived via straight forward calculations involving eigenfunctions derivatives and FPC scores pseudo inverse [26–28], offer a geometric perspective on local developmental progression and complement the global pseudotime covariance captured in the FDA model (see Supplementary Section 1.3 for details). Projecting cells into the FPC space showed coherent cluster structure and cell state transitions as expected by the biological cell developmental trajectory and pseudotime estimated by *Monocle3*, providing a strong internal validation of the useful information captured in top FPC space and the cell trajectory derived by the pseudotime analysis (Figure 4c–d).

**Fig. 4.**
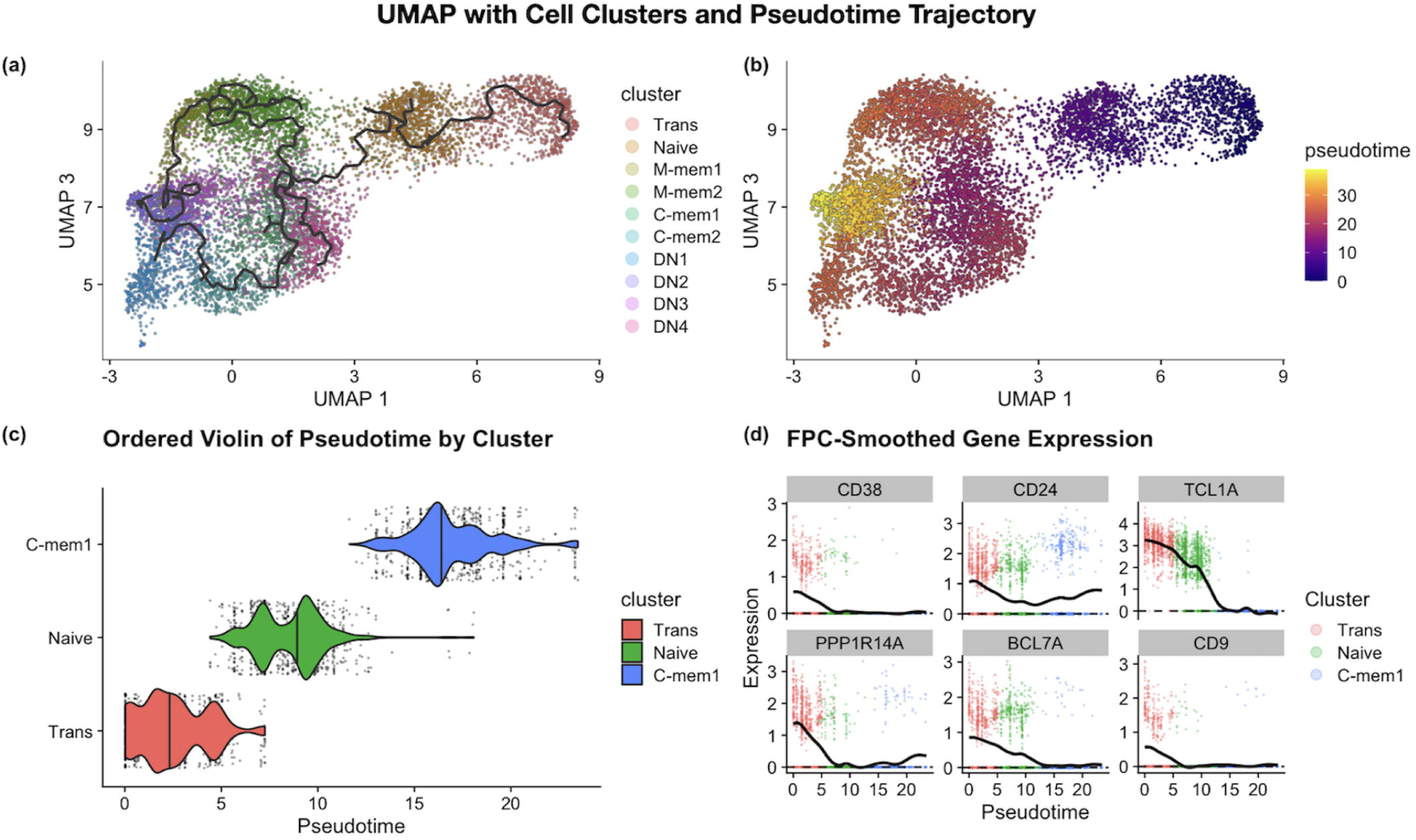
FPC space reconstruction in real data analysis. (a) Top 3 FPC eigenfunctions explain 92.6% of the total variation along the cell trajectory between Trans, Naïve, and C-mem1 clusters, representing the major pseudo-temporal gene expression patterns. (b) Pseudotime covariance contour structure shows regions of high gene expression interactions along cell pseudotime progression. (c) Projecting cells onto the FPC space deciphers the cell development trajectory, with directional vectors indicating cell direction along the cell trajectory. (d) Cells projected in the FPC space with pseudotime estimated by *Monocle3*.

We also compared the results by using scFPC-DE and PseudotimeDE in the real dataset. Out of 2980 total genes, scFPC-DE identified 474 TDEGs and PseudotimeDE identified 1795 TDEGs along the Trans, Naïve, and C-mem1 trajectory. In Figure 5a, we compare the distributions of FDR-corrected p-values obtained from scFPC-DE and PseudotimeDE. The p-value histogram from scFPC-DE shows a relatively uniform distribution with a modest concentration near zero, reflecting conservative type I error control and a more stringent DEG selection. In contrast, PseudotimeDE yields a sharp spike near zero with minimal values elsewhere, indicating potential inflation of false positives. In Figure 5b, the TDEG results are projected in the FPC score space. The scFPC-DE result led to a clear separation between null genes and TDEGs, proving a stable decision boundary. This stable separation is achieved by using the projected *L*^2^ distance (*D_i_*) as the test statistic, which quantifies the deviation from a constant mean gene expression function in FPCA. This distance measure provides a meaningful ranking of genes in the FPC score space after the effective variance decomposition through FPCA. In Figure 6, we visualized the top 10 TDEGs identified by scFPC-DE. The smoothed functions of the top TDEGs exhibit pronounced dynamic expression patterns across pseudotime, including the predicted downregulation of IgD, a known hallmark of B cell maturation.

**Fig. 5.**
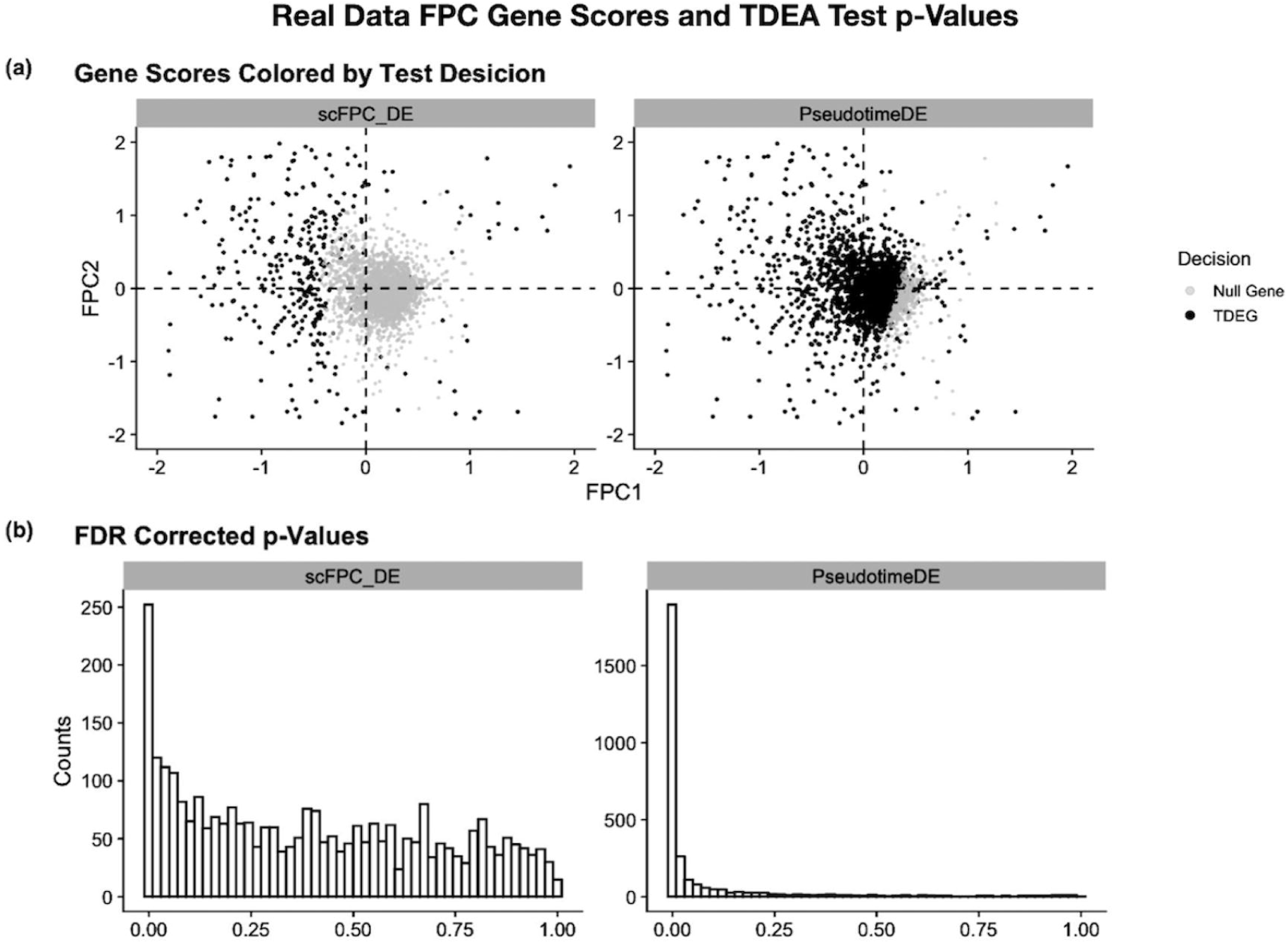
scFPC-DE and PseudotimeDE comparison on real data. (a) FPC gene score projection identifies highly differentially expressed genes, showing a well-defined boundary between the null genes and TDEGs identified by scFPC-DE. In contrast, PseudotimeDE lacks a clearly defined boundary. (b) The distribution of FDR-corrected p-values between scFPC-DE and PseudotimeDE.

**Fig. 6.**
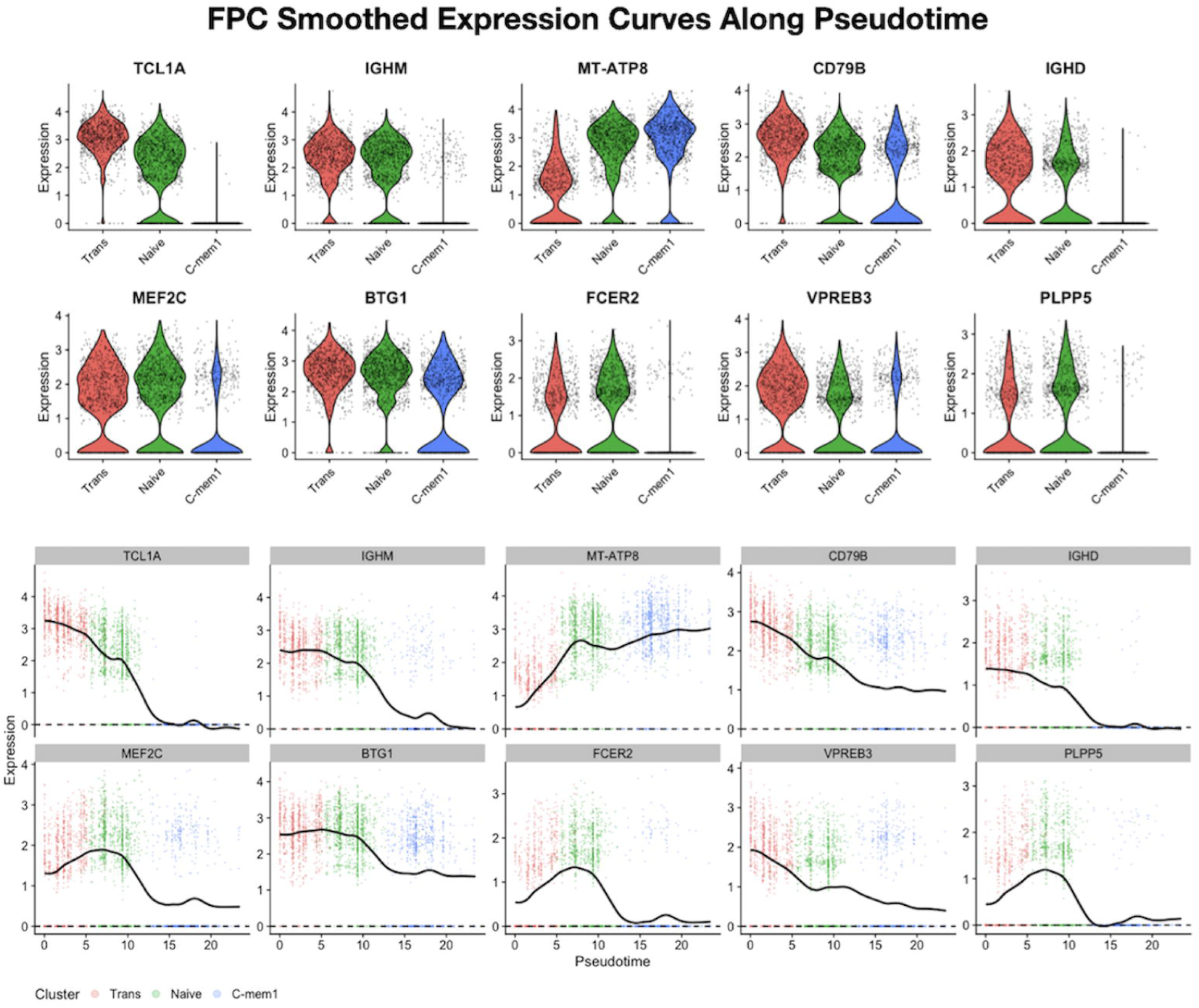
Gene expression patterns of the top TDEGs identified by scFPC-DE. Top 10 TDEGs ranked by the scFPC-DE projected distance statistic allows further inspection for TIGs along cell pseudotime progression. Violin plots grouped by the discrete clusters as well as the smoothed gene expression functions are plotted for each top TDEG. The functional data representation offers a dynamic view of the continuous gene expression changes across the cell states.

### 3.3 Pathway enrichment analysis

We used the *ReactomePA* R package [18] and the *Enrichr-KG* online tool (https://maayanlab.cloud/enrichr-kg) [19] to conduct pathway enrichment analysis for the TDEG lists identified by scFPC-DE and PseudotimeDE. In addition to the Reactome pathways, by using *Enrichr-KG*, we included more biologically curated pathway and phenotype ontologies, such as the KEGG pathways, the Mammalian Phenotype Ontology [29], the Human U133A/GNF1H Gene Atlas [30] and the HuBMAP ASCT+B tables for single cell-specific annotations [31]

Despite identifying substantially fewer TDEGs (Figure 7a), scFPC-DE uncovered 43 uniquely enriched pathways that are associated with B-cell immune functions and lineage progression in the Reactome pathway analysis (Figure 7b; Supplementary Table 4). While both scFPC-DE and PseudotimeDE identified 25 common pathways (Figure 7b), the four enriched pathways unique to PseudotimeDE reflected broad cellular functions with limited immune relevance (Supplementary Table 5). This contrast indicates higher biological specificity of the scFPC-DE result by precisely modeling the functionally coherent gene programs and avoiding false discoveries. These results are consistent with our simulation observations, where scFPC-DE maintained sufficient power while substantially reducing false positives, especially for data with high zero inflation. Both methods recovered infection-related pathways (Supplementary Table 6) in which B cells play important roles, however scFPC-DE achieved this with only a quarter of the input TDEGs compared with PseudotimeDE. Notably most immune-specific pathways were uniquely identified by scFPC-DE. This demonstrates that scFPC-DE can identify a more compact set of TDEGs with higher signal-to-noise ratio and biologically more coherent. We additionally examined the 25 Reactome pathways shared across both methods. Enrichment p-values were consistently and substantially smaller when using the scFPC-DE gene set, reflecting stronger statistical support for true pathway-level associations. Using Over-Representation Analysis (ORA) summary statistics from *ReactomePA*, the most striking result highlighting the advantage of scFPC-DE is portrayed in Figure 7c, which compares the enrichment strength between scFPC-DE and PseudotimeDE using gene-to-background ratio plots. scFPC-DE identified immune and developmental pathways with higher gene ratios and greater statistical significance, while PseudotimeDE primarily highlighted broad categories such as mRNA splicing with weaker enrichment signals. Notably, the largest gene-to-background ratio was observed for the Immune System pathway, reinforcing the biological context of this analysis and underscoring the relevance of scFPC-DE in capturing immune-specific gene programs. Overall, scFPC-DE yields consistently stronger pathway enrichment signals in both effect size and statistical significance compared to PseudotimeDE. In summary, scFPC-DE identifies a more concise and biologically informative set of TDEGs for pathway analysis.

**Fig. 7.**
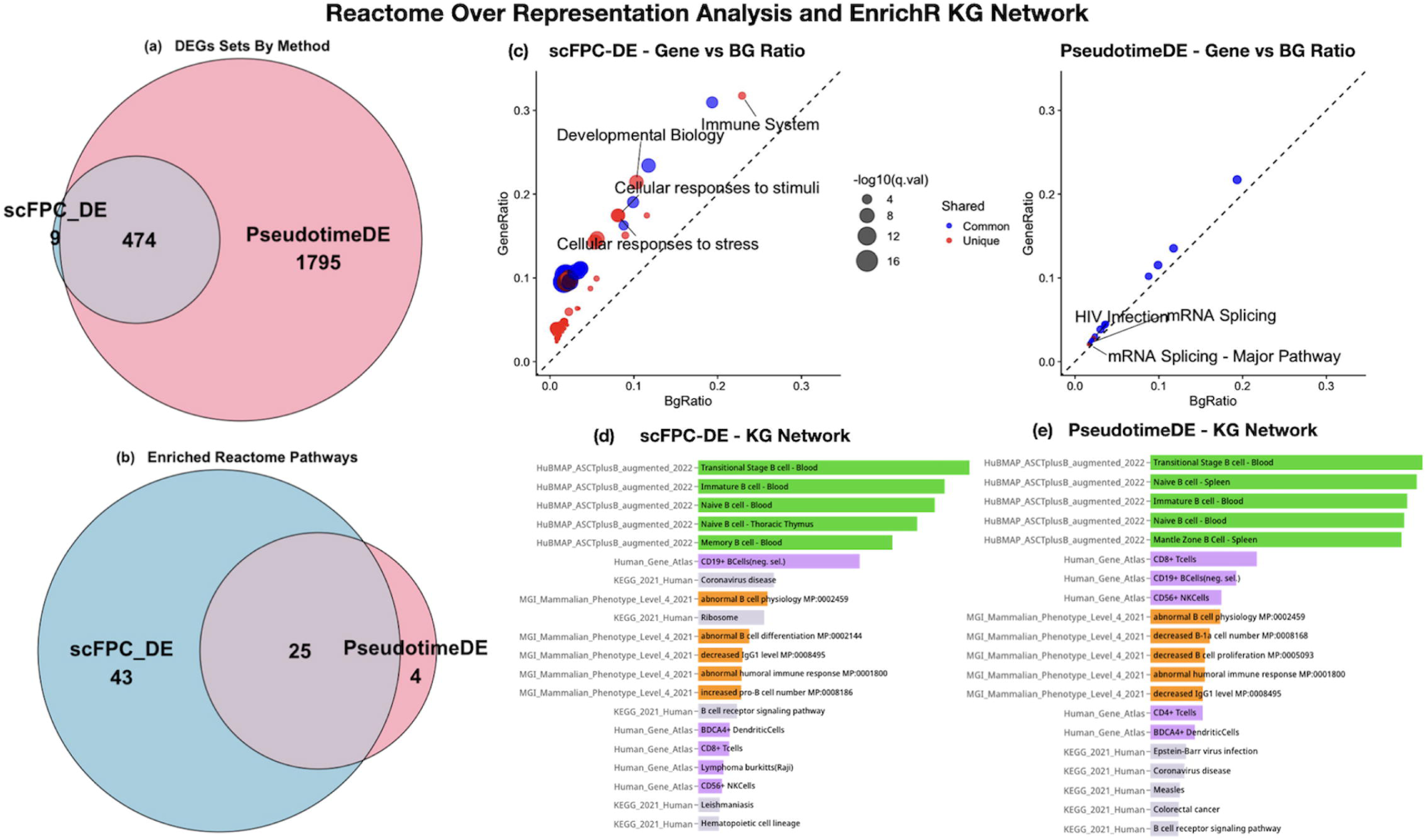
Pathway and phenotype-level enrichment analysis results for scFPC-DE and PseudotimeDE. (a) Number of method-specific and common TDEGs identified by scFPC-DE and PseudotimeDE. scFPC-DE identified a smaller, more targeted set of TDEGs—approximately one-fourth the size of that returned by PseudotimeDE. (b) Number of method-specific and common pathways identified in the Reactome pathway analysis. scFPC-DE uniquely identified 43 pathways, PseudotimeDE identified 4 unique pathways, and 25 pathways were shared between the two methods. Even when sharing most of the same DEGs, PseudotimeDE did not identify these unique pathways when following the same GSEA protocol. (c) The gene-to-background ratio plots show that the enrichment signals from scFPC-DE are stronger for both shared and unique pathways. (d–e) The *Enrichr-KG* results for pathway- and phenotype-level enrichment analysis using multiple ontologies (MGI Mammalian Phenotype Level 4, HuBMAP ASCT+B 2022, Human Gene Atlas, and KEGG 2021).

To further assess the biological relevance of the TDEG lists from the two DE methods, we used the convenient *Enrichr-KG* online tool to query more biological entity databases. Figure 7d-e show the top 5 terms from the selected ontologies. Using the single cell-specific knowledge curated in the HuBMAP ASCT+B tables (green), the scFPC-DE list correctly annotated the B cells from blood as PBMC samples were used in the original study (Stewart, et al., 2021). The thoracic thymus is a critical hub for developing central immune tolerance; thus, it would be expected to contain similar populations of Naïve B cell. On the contrary, the PseudotimeDE list annotated two B cell populations from spleen – an irrelevant organ to the real data, therefore the TDEG list is likely to contain misinformation from false discoveries. Both methods recognized specific terms related to developmental and mature B cell phenotypes and functions, such as the CD19+ B cells (purple), humoral immune response (orange), and B cell receptor signaling pathway (grey), with higher ranking in the scFPC-DE output ordered by the combined score of *Enrichr-KG* than in the PseudotimeDE output. Supplementary Figure 9 also shows the knowledge graph visualization of biological phenotypes and connected genes derived from the TDEG lists, revealing gene-specific contributions to the phenotypes.

Together, these results illustrate that scFPC-DE can effectively detect a more interpretable, statistically robust, and biologically meaningful enrichment profile than existing method such as PseudotimeDE.

## Discussion

In this study, we introduced scFPC-DE, a novel method based on functional data analysis techniques for trajectory-based differential expression analysis using scRNA-seq data. By leveraging FPCA, scFPC-DE provides a continuous, data-driven representation of gene expression dynamics along the cell trajectory reconstructed by pseudotime. The FPCA modeling framework captures the dominant variation across genes while preserving the temporal pattern inherent to the underlying cellular progression in the cell trajectory, enabling a more integrated and interpretable model of gene expression changes.

Simulation studies, designed to reflect the high zero inflation phenomenon in the real scRNA-seq data, demonstrate that scFPC-DE offers substantial advantages over existing approaches. It consistently achieved the best control of type I error while attaining the highest AUC in the ROC analysis. The improved overall performance is largely attributed from its ability to borrow information across genes and across pseudotime, in contrast to the GAM-based methods that model each gene independently and lack global covariance modeling.

While PseudotimeDE and other GAM-based approaches have advanced the modeling of nonlinear trajectories, they fail to incorporate inter-gene dependencies. In contrast, scFPC-DE’s FPCA-based approach effectively captures the global inter-gene variance-covariance structure in the top FPC space and uses functional projection to distinguish null from alternative hypotheses in the FPC space with a robust null boundary that leads to a tighter control of type I error, particularly for scRNA-seq data with high zero inflation. Application to a well-characterized B cell subtypes dataset further illustrates the advantages of scFPC-DE. With a more concise list of DEGs identified by scFPC-DE than by PseudotimeDE, the scFPC-DE identified DEG list revealed specific immune pathways related to B cell functions in Reactome enrichment analysis, indicating a stringent control of false positive DEGs by scFPC-DE. The real data analysis suggests that scFPC-DE can effectively avoid those noises associated with a large number of false positive genes, yielding a cleaner and more biologically coherent enrichment analysis.

One limitation in the current scFPC-DE pipeline is that it takes in the estimated pseudotime as input data, which may potentially impact the inference. In this study, our results indicate that our FPCA-based framework remains robust to variations in the input pseudotime, as shown by its consistent performance across simulations and real data. As a natural extension, we plan to develop an end-to-end FDA-based pipeline that integrates pseudotime inference and differential expression testing into a unified functional framework. We also plan to extend scFPC-DE to support complex branching trajectories and to include functional pseudotime estimation using the same FDA framework.

In conclusion, scFPC-DE defines a statistically rigorous and biologically interpretable framework for trajectory-based differential expression analysis. Its ability to model smooth gene expression variation across pseudotime, incorporate inter-gene covariance, and control false discoveries marks a significant advancement over existing methods.

## R Package Availability

The scFPC-DE R package implementing all methods described in this study is publicly available at: https://github.com/LopezRicardo1/scFPCDE.

