## Supplementary Material for "scFPC-DE: Robust Differential Expression Analysis Along Single Cell Trajectories via Functional Principal Component Analysis"

Supplementary Text

Ricardo J. López Candelaria[1], Yu Qian[2], Fang Chen[3], Mansun Law[3], Yun Zhang[4*], and Xing Qiu[1*]

**4 References** **17**

[1] Department of Biostatistics and Computational Biology, University of Rochester, NY, USA
[2] Department of Informatics, J. Craig Venter Institute, La Jolla, CA, USA
[3] Department of Immunology and Microbiology, Scripps Research, La Jolla, CA, USA
[4] Division of Intramural Research, National Library of Medicine, NIH, Bethesda, MD, USA

*To whom correspondence should be addressed.
Contact:

### 1 Trajectory Representation in FPC Space

#### 1.1 Covariance Operator from Pseudotime-Centered Functional Gene Profiles

To characterize shared variation across genes along pseudotime, we begin by constructing the empirical covariance operator from smooth gene expression curves. Each gene is treated as a function $y_j(t)$ of pseudotime $t \in [0,1]$, and centered in the pseudotime direction:

$$\tilde{y}_j(t) = y_j(t) - \frac{1}{T}\sum_{t=1}^{T} y_j(t) \tag{1}$$

This removes the average trajectory over pseudotime for each gene, emphasizing deviation patterns that are temporally structured.

The empirical covariance operator $C(s,t)$ is then computed by averaging across all genes:

$$C(s,t) = \frac{1}{p}\sum_{j=1}^{p} \tilde{y}_j(s)\tilde{y}_j(t) \tag{2}$$

This operator captures the pseudotime-dependent structure in the data and provides a smooth, low-rank representation of gene co-variation.

As referred in equation 2 from the main text, the dominant temporal patterns satisfy following integral equation:

$$\int C(s,t)\Phi_k(s)\,ds = \lambda_k \Phi_k(t) \tag{3}$$

where $\Phi_k(t)$ are the orthonormal eigenfunctions and $\lambda_k$ the associated eigenvalues. The full covariance can then be expressed as a spectral decomposition:

$$C(s,t) = \sum_{k=1}^{\infty} \lambda_k \Phi_k(s)\Phi_k(t) \tag{4}$$

This decomposition is automatically performed in the scFPC-DE pipeline using standard methods from the fda package (J. O. Ramsay and Silverman 2005), with a slight modification in the data pre-processing step to ensure pseudotime-centered input functions.

After selecting the top $L$ eigenfunctions (e.g., explaining a pre-specified proportion of total variance), we approximate the covariance structure by:

$$\hat{C}(s,t) = \sum_{k=1}^{L} \lambda_k \Phi_k(s)\Phi_k(t) \tag{5}$$

The smooth low-rank structure of $\hat{C}(s,t)$ enables visualization of local developmental cell pseudotime interactions associated with high gene expression. In the main text (Figure 4b), we display the reconstructed pseudotime covariance structure as contour plot.

#### 1.2 Functional Expansion and Score Space Projection

As described in Section 3.3 of the main manuscript, each gene's smoothed expression trajectory across cells is represented using a functional principal component expansion over eigenfunctions $\{\phi_k(t)\}_{k=1}^{L}$, derived from FPCA (Jolliffe and Cadima 2016; Shang 2014):

$$y_i(t_j) = \sum_{k=1}^{L} \xi_{ik}\, \phi_k(t_j) + \varepsilon(t_j), \quad j = 1, \dots, T, \tag{6}$$

where $\phi_k(t_j)$ is the $k$-th eigenfunction evaluated at pseudotime $t_j$, and $\xi_{ik}$ is the corresponding FPC score for gene $i$. The full score matrix $\boldsymbol{\xi} \in \mathbb{R}^{G \times L}$ contains rows $\boldsymbol{\xi}_i$ for each gene $i = 1, \dots, G$.

The FPC scores are computed via regression. Let $\hat{\mathbf{y}}_i = (\hat{y}_i(t_1), \dots, \hat{y}_i(t_T))^\top \in \mathbb{R}^T$, and let $\boldsymbol{\Phi} \in \mathbb{R}^{T \times L}$ be the matrix whose columns are the eigenfunctions evaluated at each pseudotime point. Then the scores $\boldsymbol{\xi}_i \in \mathbb{R}^L$ are estimated by:

$$\hat{\boldsymbol{\xi}}_i = \left( \boldsymbol{\Phi}^\top \boldsymbol{\Phi} \right)^{-1} \boldsymbol{\Phi}^\top \hat{\mathbf{y}}_i. \tag{7}$$

Once estimated, we can project the observed expression data $\mathbf{Y} \in \mathbb{R}^{G \times T}$ into the FPC score space using the pseudo-inverse (MacAusland 2014; Prasad and Bapat 1992) of the score matrix, $\mathbf{Y}\hat{\boldsymbol{\xi}}^{+}$, which maps each cell to a location along the trajectory defined by the eigenbasis.

#### 1.3 Derivative-Based Vector Field Construction

To visualize pseudotime orientation in the FPC score space, we compute the derivatives of the eigenfunctions:

$$\phi_k'(t) = \frac{d}{dt}\phi_k(t), \quad k = 1, \dots, L. \tag{8}$$

These derivatives define directional vectors (gradient field):

$$\mathbf{v}(t) = \left(\phi_1'(t), \dots, \phi_L'(t)\right)^\top, \tag{9}$$

which encode the local direction of progression along the FPC space.

In real data applications, it is often sufficient to use the first two components to project the direction vector, represented as arrows, into a 2D plane:

$$\mathbf{v}_j^{(2)} = \left(\phi_1'(t_j),\ \phi_2'(t_j)\right)^\top, \tag{10}$$

which defines the orientation of progression at cell $j$ in the reduced FPC space.

These directional vectors are shown in Figure 4 of the main manuscript. All quantities above are computed using the `fda` framework (J. Ramsay et al. 2009).

### 2 Simulation Framework and Zero-Inflation Sensitivity

#### 2.1 Simulation Design Based on scDesign3

We used the `scDesign3` framework to simulate realistic single-cell expression data guided by B-cell differentiation trajectories, as described in Section 3.1 of the main manuscript (Song et al. 2024). The generative model followed a zero-inflated negative binomial (ZINB) distribution to capture both biological variability and technical dropout.

To introduce increasing levels of sparsity, we applied a Bernoulli masking process (Equation 5 main text) to the simulated counts with the following dropout probabilities:

- ZI0: no additional dropout (baseline)
- ZI1: Bernoulli dropout probability = 0.25
- ZI2: Bernoulli dropout probability = 0.50
- ZI3: Bernoulli dropout probability = 0.75

Each dataset includes 4,000 genes measured across 500 cells, all sharing a common pseudotime progression. The first 500 genes were designated as trajectory differentially expressed (TDEG), with the remaining 3,500 serving as null genes. This setup enables controlled evaluation of both Type I error and statistical power under a known ground truth.

The following table summarize the zero proportions and descriptive statistics of log-transformed data sets for each zero-inflation condition.

Table 1: Summary of zero proportion and log transformed count distribution across the four simulated datasets (ZI0–ZI3).

| Dataset | Zero_Proportion | Min | Q1 | Median | Q3 | Max | Mean | SD |
|---|---|---|---|---|---|---|---|---|
| ZI0 | 0.647 | 0 | 0 | 0 | 1 | 8.24 | 0.49 | 0.75 |
| ZI1 | 0.717 | 0 | 0 | 0 | 1 | 8.52 | 0.39 | 0.69 |
| ZI2 | 0.823 | 0 | 0 | 0 | 0 | 9.45 | 0.24 | 0.58 |
| ZI3 | 0.931 | 0 | 0 | 0 | 0 | 7.36 | 0.10 | 0.38 |

All datasets shared the same underlying gene expression dynamics and oracle pseudotime, enabling controlled benchmarking across conditions. As a remark, we point out that the no-dropout-condition sparsity percentage reflects the true biological zeros—that is, instances where genes are genuinely not expressed in specific cells (Jiang et al. 2022). This baseline offers a reference level of sparsity before introducing additional dropout.

#### 2.2 Benchmarking Under Varying Zero-Inflation Levels

To evaluate the robustness of differential expression (DE) methods under varying dropout scenarios, we conducted a comprehensive sensitivity analysis across the four simulated datasets (ZI0–ZI3), each corresponding to an increasing level of zero inflation. The simulation study used the oracle pseudotime—that is, the true latent ordering used to generate the data—allowing us to assess the statistical inference properties of each method without the confounding effects of trajectory estimation (Song et al. 2024).

We applied five DE methods to each dataset:

- scFPC-DE (FPC $L^2$ distance-based)
- FPC-F (F-statistic-based)

- PseudotimeDE (negative binomial model)
- PseudotimeDE_ZiNB (zero-inflated negative binomial model)
- PseudotimeDE_G (Gaussian model)

The performance of each method was assessed using multiple criteria:

- Type I error control: comparing observed false positive rates to expected values across significance levels
- Statistical power: proportion of true DE genes detected at fixed thresholds
- Area under the ROC curve (AUC): measuring overall ranking performance
- FPC score space projections: visualizing separation of DE vs. non-DE genes
- Computation time: efficiency benchmarking across zero-inflation levels

scFPC-DE demonstrated consistent performance across increasing dropout levels, maintaining both type I error control and statistical power. In contrast, PseudotimeDE [RN29], using a ZINB model exhibited increased computation time and reduced stability under the highest dropout condition (ZI3), underscoring the sensitivity of its inference procedure to zero inflation and model complexity.

As a remark, we note that the ZI3 condition closely mirrors the zero-inflation profile observed in the real dataset analyzed in Section 3.2 of the main manuscript. This alignment was intentional, allowing the ZI3 simulation to benchmark the robustness of scFPC-DE under conditions that realistically reflect biological single-cell data.

The following results summarize the reconstructed gene expression curves from simulated datasets that emulate the sparsity and zero-inflation patterns characteristic of real scRNA-seq data. Figure 1 illustrates the expression-generating functions described in Equation 5, alongside smoothed gene expression trajectories over oracle pseudotime. These trajectories result from the combined contributions of the mean function and stochastic variation as defined in Equation 6 of the main text. Top-ranked DE genes exhibit dynamic expression patterns over pseudotime, while null genes display flat profiles.

In Figure 2, the AUC metric degrades as zero inflation increases, but scFPC-DE consistently achieves the highest AUC across all dropout levels. Figures 3 and 4 show that scFPC-DE maintains accurate Type I error control and attains the highest statistical power, especially under high dropout scenarios. Figure 5 illustrates the FPC score projections, highlighting that scFPC-DE preserves a well-defined null boundary even under severe zero inflation, in contrast to other methods whose null regions become diffuse. Finally, Figure 6 and Supplementary Table 2 show that scFPC-DE exhibits stable runtimes across dropout conditions and is more than 10 times faster on average than PseudotimeDE, particularly under the ZINB model (Song and Li 2021; Song et al. 2024; Jiang et al. 2022).

Table 2: Summary stats in seconds across all zero inflation levels by method

| Method | n | mean | median | sd | min | max | iqr |
|---|---|---|---|---|---|---|---|
| scFPCDE | 4 | 4.91 | 4.93 | 0.30 | 4.53 | 5.26 | 0.28 |
| PseudotimeDE_G | 4 | 12.89 | 11.23 | 3.53 | 10.95 | 18.17 | 2.22 |
| PseudotimeDE | 4 | 62.80 | 48.08 | 52.72 | 16.70 | 138.35 | 39.43 |
| PseudotimeDE_ZINB | 4 | 1763.73 | 2122.84 | 911.87 | 409.96 | 2399.26 | 499.23 |

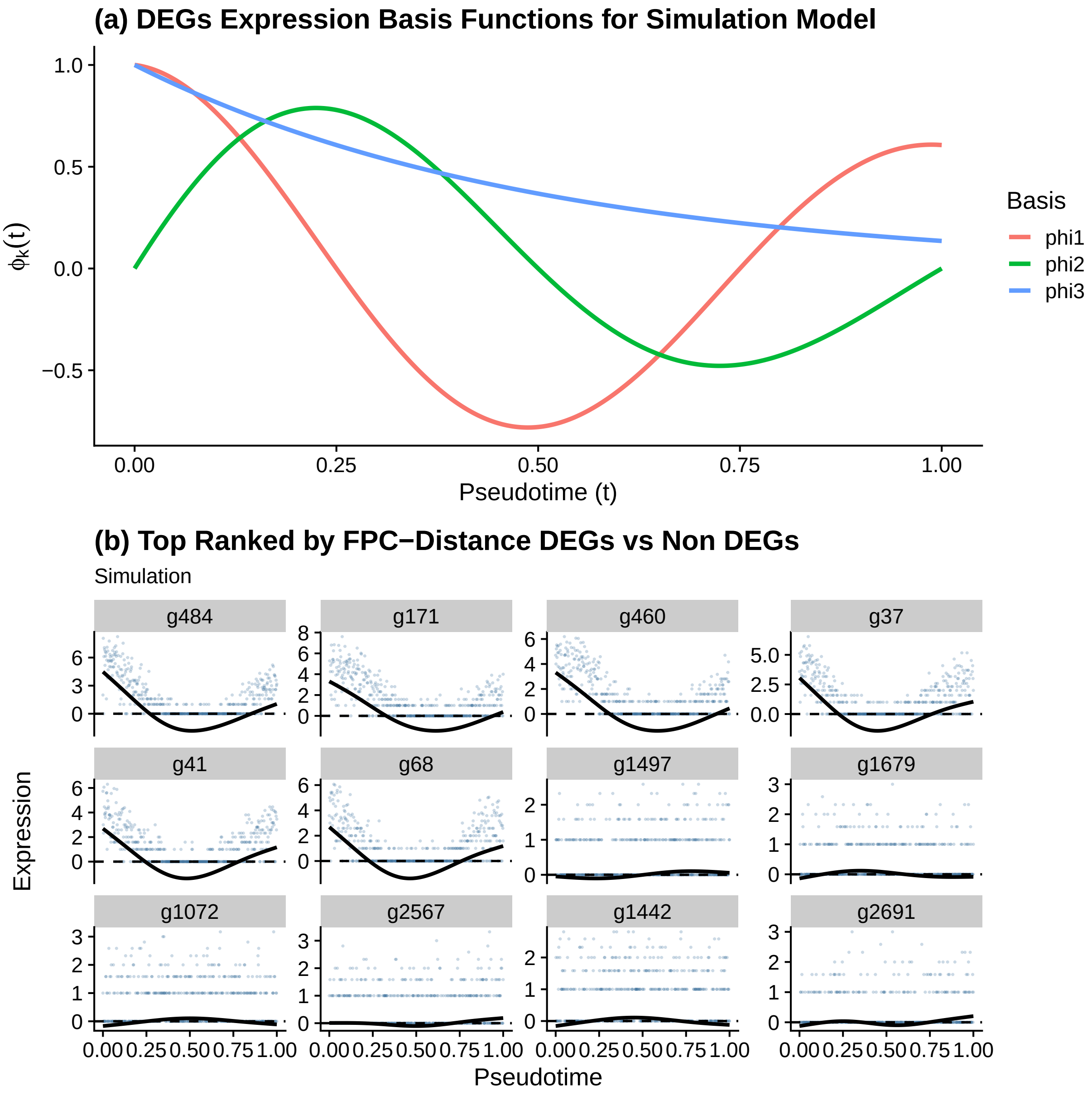

Figure 1: (a) DEGs expression basis functions used in the simulation model to define pseudotime-dependent gene expression. Each simulated gene is a linear combination of these smooth basis curves. (b) Top 15 genes ranked by the FPC-distance test statistic from the simulated dataset. Each panel shows gene expression versus pseudotime with fitted smoothed expression curves. Genes g484, g171, g460, and others display clear dynamic trends, highlighting the ability of scFPC-DE to separate dynamic from non-dynamic expression patterns.

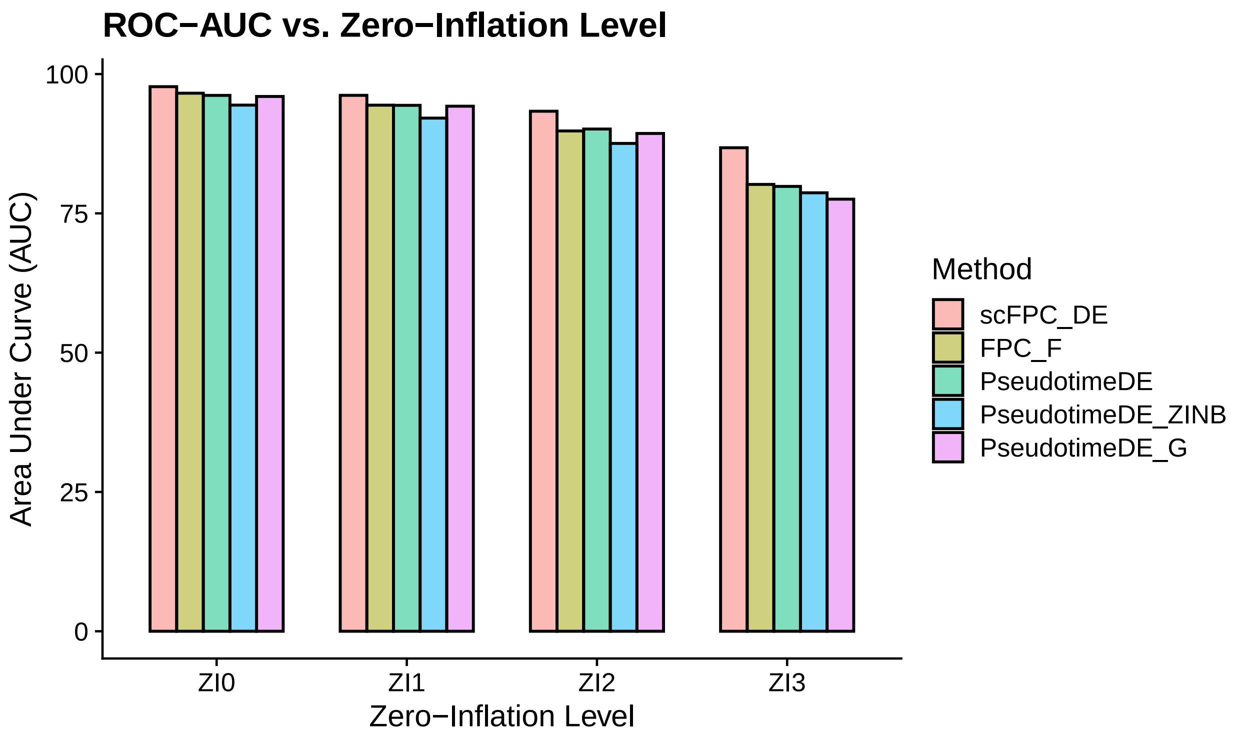

Figure 2: **Area Under the ROC Curve (AUC) across zero-inflation levels.** Each line tracks AUC performance of a method (scFPC_DE, FPC_F, PseudotimeDE, PseudotimeDE_ZINB, PseudotimeDE_G) across increasing levels of zero inflation (ZI0 to ZI3). scFPC_DE consistently achieves the highest AUC across all settings.

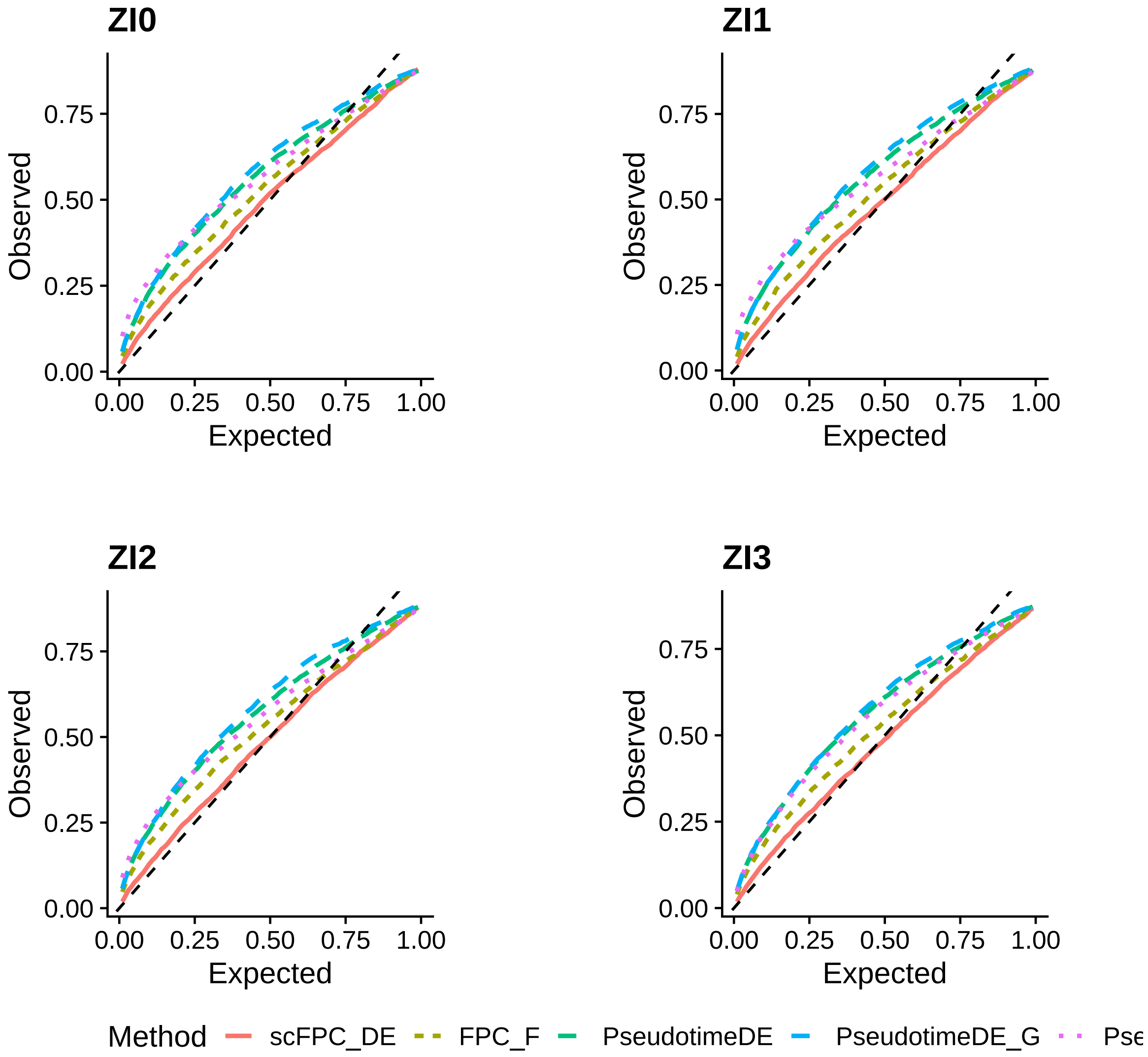

Figure 3: **Type I Error Curves across zero-inflation levels.** Each panel shows the observed Type I error rate versus nominal significance level for different methods under a given zero-inflation setting (ZI0–ZI3). scFPC_DE consistently aligns more closely to the diagonal, indicating stronger control of false positives.

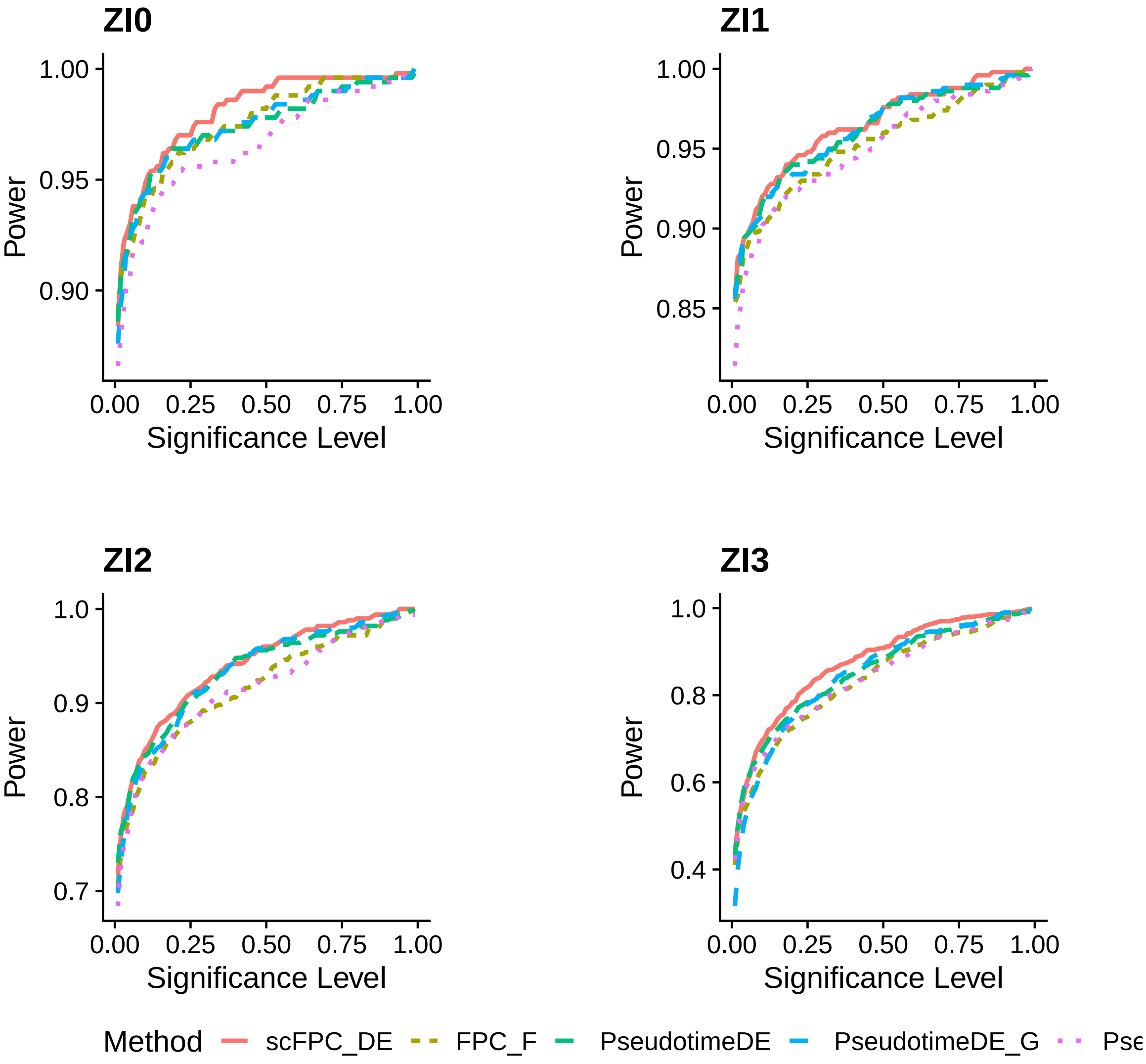

Figure 4: **Power Curves across zero-inflation levels.** Each panel displays the empirical power curves of scFPC_DE and competing methods under increasing levels of zero-inflation (ZI0–ZI3). scFPC_DE consistently demonstrates higher power, especially in more challenging scenarios.

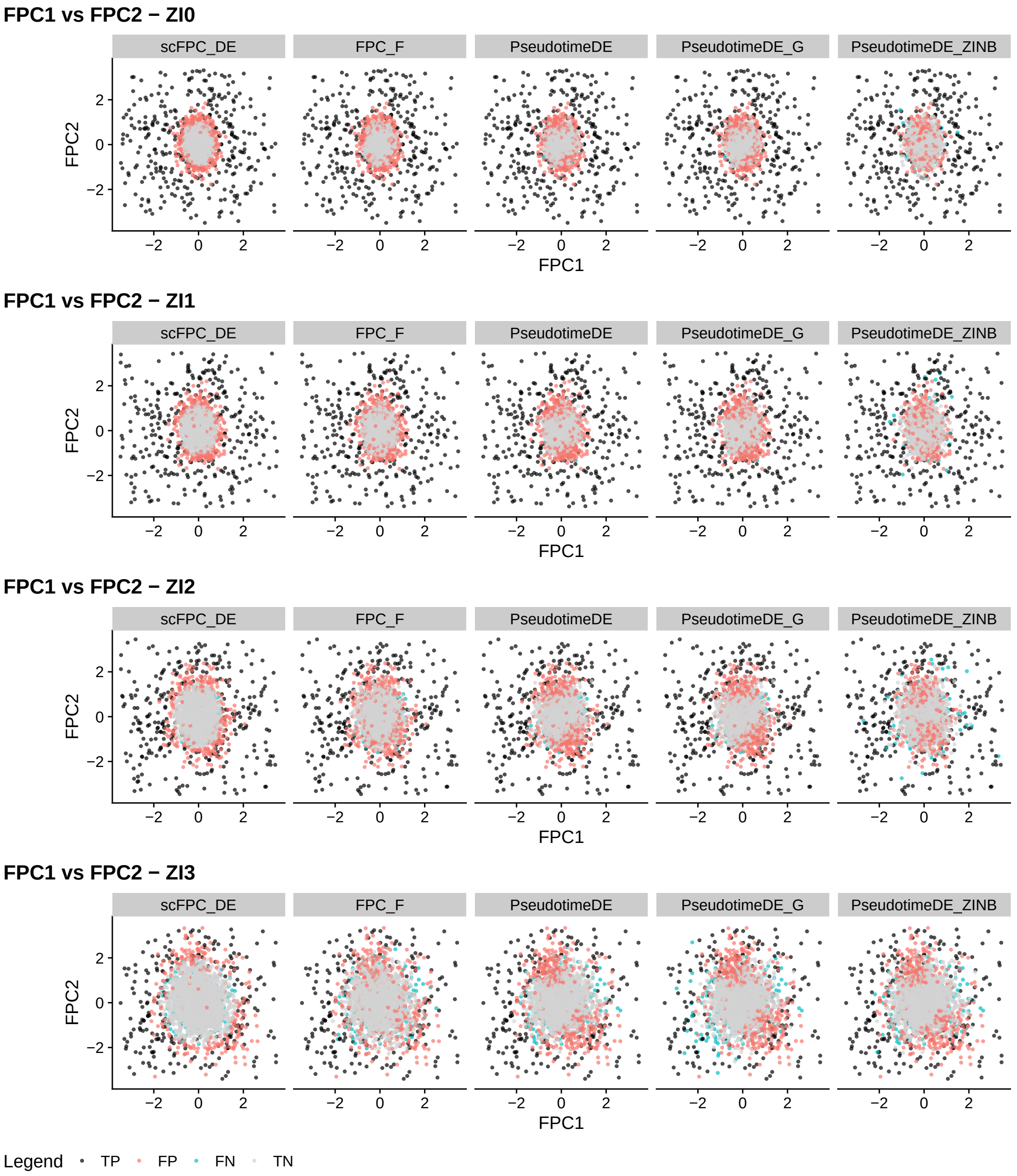

Figure 5: **Gene FPC-score space across zero-inflation levels**. Rows correspond to ZI0–ZI3. Within each row, the five columns show results for scFPC_DE, FPC_F, PseudotimeDE, PseudotimeDE_ZINB, and PseudotimeDE_G. Points are coloured by outcome: true positives (black), false positives (red), false negatives (blue), and true negatives (gray). scFPC_DE consistently recovers a well-defined null region across all zero-inflation levels, demonstrating stability of its decision boundary despite increasing dropout.

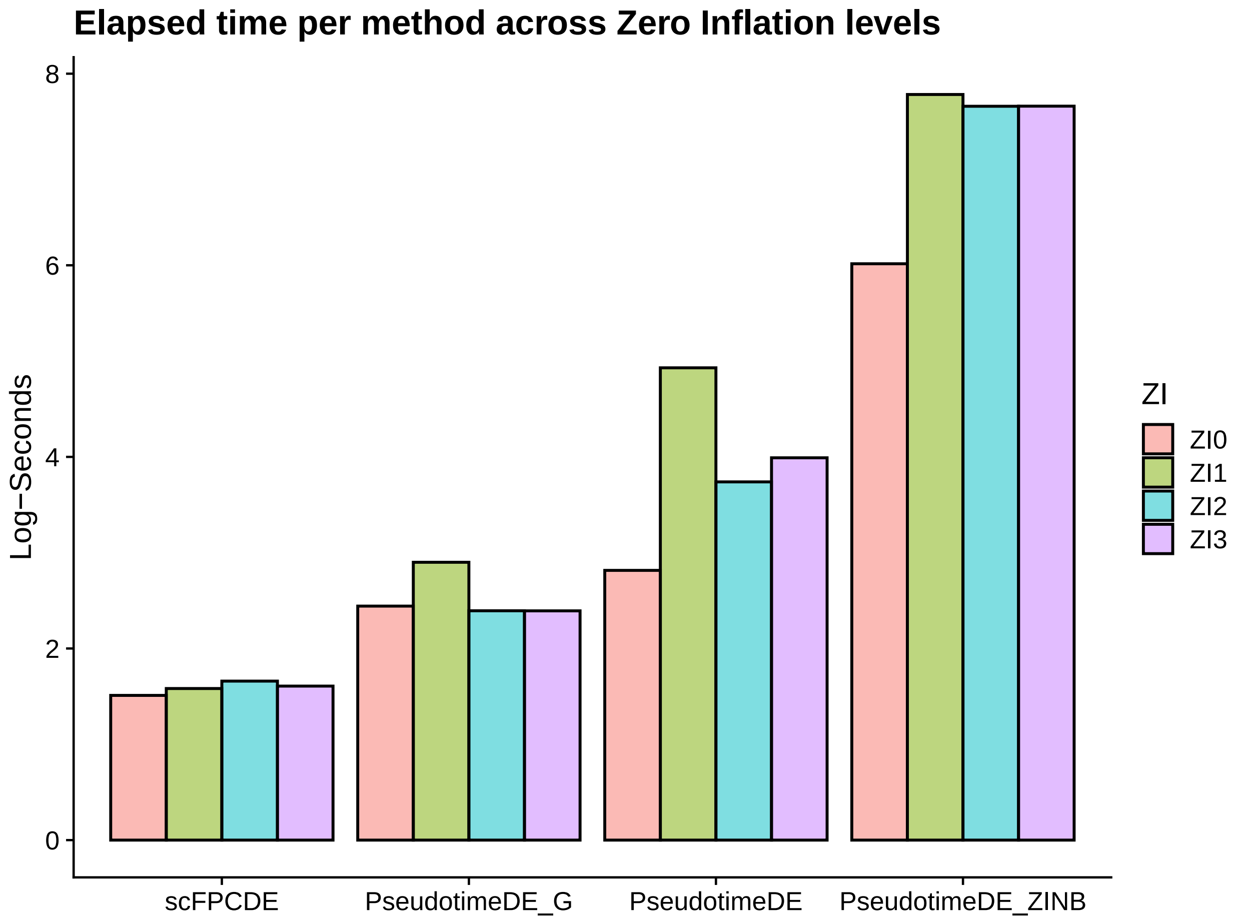

Figure 6: **Computation time across zero-inflation scenarios**. Bar plot showing the natural logarithm of the elapsed time (in seconds) for each differential expression method—scFPC-DE, PseudotimeDE-G (Gaussian), PseudotimeDE (NB), and PseudotimeDE-ZINB—across four levels of zero inflation (ZI0 to ZI3). scFPC-DE maintains stable runtimes regardless of zero inflation, while PseudotimeDE-ZINB shows increasing computation time as zero inflation increases.

### 3 Real Data Application: B Cell Trajectory

We performed an additional trajectory-aware filtering step beyond standard preprocessing to remove genes with predominantly zero counts across pseudotime. These genes lack sufficient temporal signal and cannot be modeled reliably using smooth functional trajectories. As shown in Figure 7, we illustrate—by example—three representative genes from the B-cell dataset (Stewart et al. 2021),highlighting the rationale for exclusion: IGKVD2-30 exhibits near-zero expression and is removed, whereas RPS27 and IGHD retain meaningful variation and are retained. Table 3 summarizes the zero proportion and distributional statistics for the final filtered dataset.

#### 3.1 Gene Filtering Based on Pseudotime Coverage

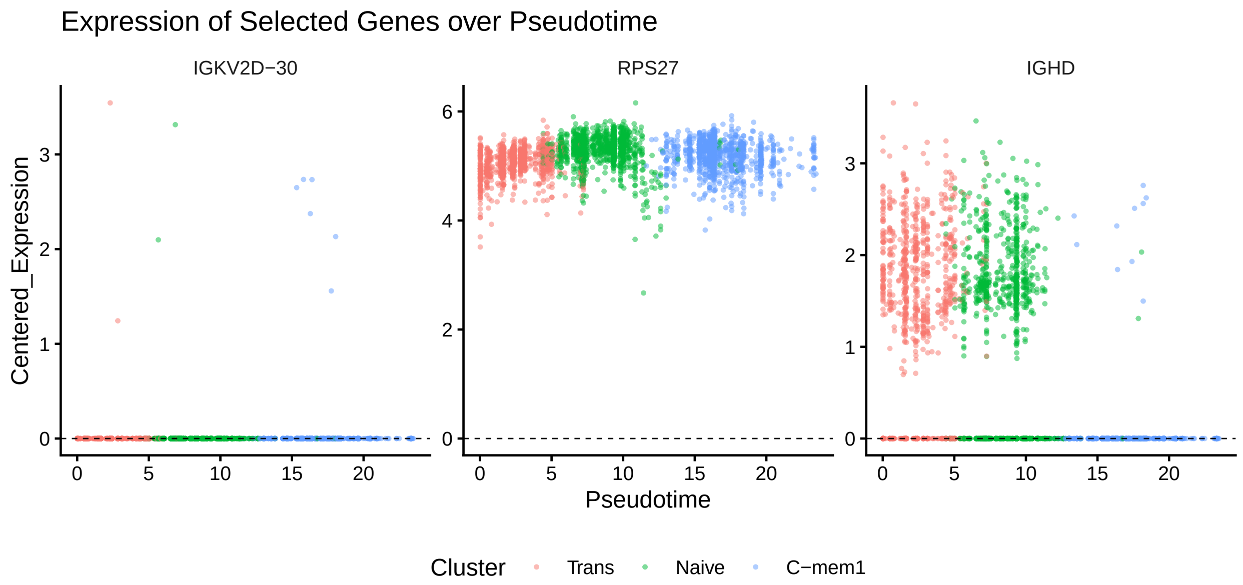

Figure 7: Expression of three genes across pseudotime; IGKVD2-30 shows near-zero coverage and is a candidate for additional pre-filtering, whereas RPS27 and IGHD remain informative along the trajectory.

Table 3: Summary of overall zero proportion and log-counts distribution for the real dataset.

| Dataset | N_Genes | n_cells | Zero_Proportion | Min | Q1 | Median | Q3 | Max | Mean | SD |
|---|---|---|---|---|---|---|---|---|---|---|
| Real-B-cell | 2980 | 2988 | 0.939 | 0 | 0 | 0 | 0 | 6.97 | 0.14 | 0.61 |

#### 3.2 Impact on FPCA Smoothing and Eigenstructure

To stabilize functional smoothing and avoid over-penalization due to high sparsity, we reconstructed the FPCA basis using only a subset variable genes with respect to the cell trajectory. Figure 8 illustrates the effect of this selection on the GCV criterion and the estimated eigenfunctions: while using all genes yields flatter curves due to excessive zero inflation, restricting to high-variability genes produces sharper temporal modes and preserves comparable variance explained. This balance improves trajectory modeling without sacrificing interpretability.

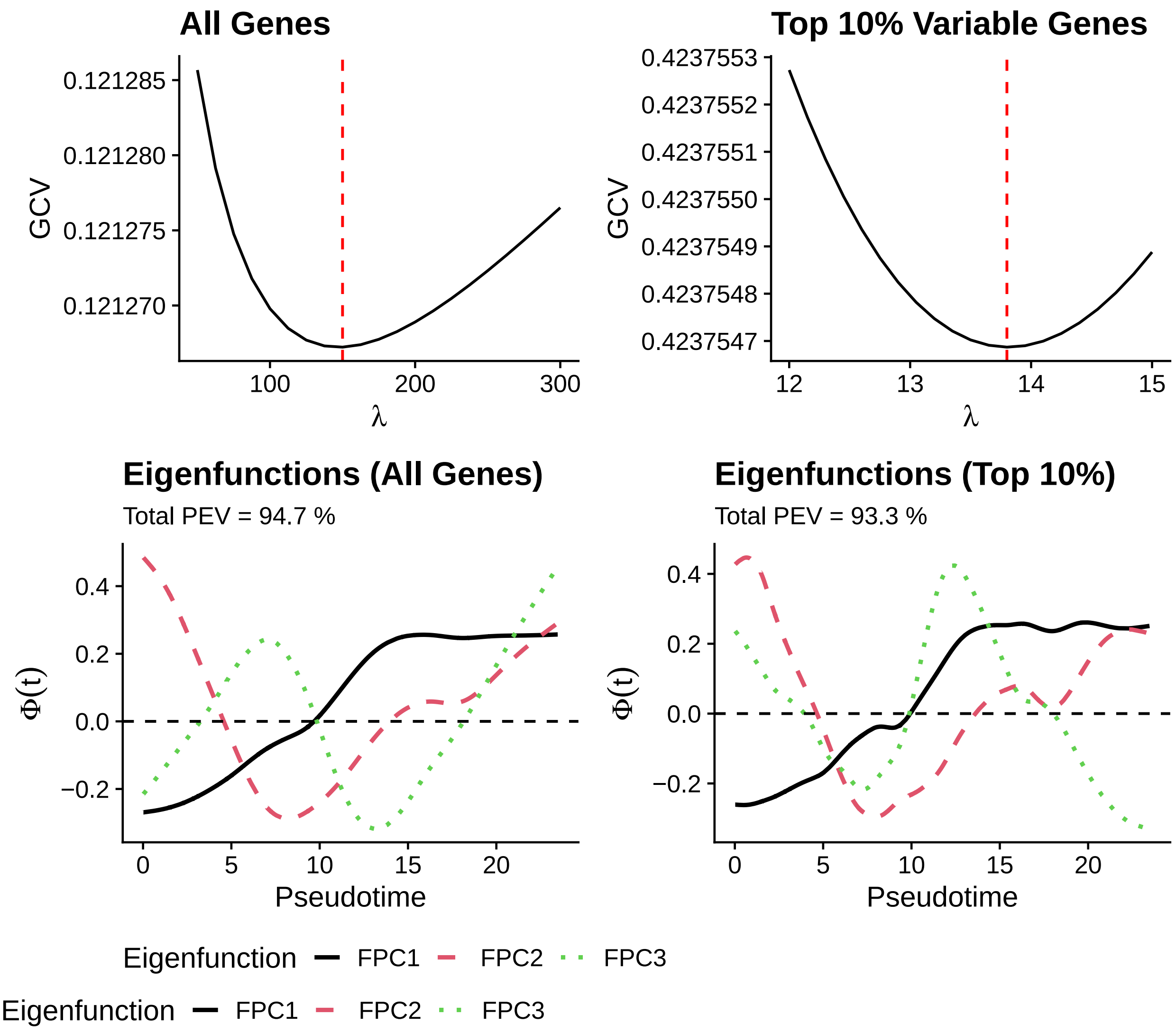

Figure 8: Influence of gene selection on FPCA smoothing and eigenfunction structure. GCV curves (top) show that including all genes biases the criterion toward higher smoothing due to zero inflation and low variability, while restricting to the top 10% most variable genes leads to more moderate penalization. Eigenfunctions (bottom) are flatter when using all genes, whereas sharper temporal modes are preserved with top-variable genes, maintaining comparable total variance explained.

#### 3.3 Reactome ORA Pathways Results

In the main text (Figure 7b), we summarized gene set over-representation analysis using ReactomePA (Yu and He 2016), comparing pathways uniquely identified by scFPC-DE and PseudotimeDE (Song and Li 2021). This comparison was based on DEGs derived from the B cell dataset described earlier. The following tables list the full set of Reactome pathways recovered by each method, categorized as unique or shared, and were used to generate the gene ratio versus background ratio visualization in Figure 7c.

Table 4: Pathways uniquely identified by scFPC-DE

| Pathway |
|---|
| Nonsense-Mediated Decay (NMD) |
| Nonsense Mediated Decay (NMD) enhanced by the Exon Junction Complex (EJC) |
| Regulation of expression of SLITs and ROBOs |
| Cellular response to starvation |
| Signaling by ROBO receptors |
| Axon guidance |
| Nervous system development |
| Developmental Biology |
| Cellular responses to stress |
| Cellular responses to stimuli |
| SARS-CoV-1 modulates host translation machinery |
| Translation initiation complex formation |
| Activation of the mRNA upon binding of the cap-binding complex and eIFs, and subsequent binding to 43S |
| Formation of the ternary complex, and subsequently, the 43S complex |
| Ribosomal scanning and start codon recognition |
| SARS-CoV-2 modulates host translation machinery |
| Fcgamma receptor (FCGR) dependent phagocytosis |
| SARS-CoV-1 Infection |
| SARS-CoV-1-host interactions |
| Immune System |
| Immunoregulatory interactions between a Lymphoid and a non-Lymphoid cell |
| Cytokine Signaling in Immune system |
| Leishmania infection |
| Parasitic Infection Pathways |
| Innate Immune System |
| SARS-CoV Infections |
| Fc epsilon receptor (FCERI) signaling |
| Signaling by the B Cell Receptor (BCR) |
| Parasite infection |
| Leishmania phagocytosis |
| FCGR3A-mediated phagocytosis |
| Hemostasis |
| GPCR downstream signalling |
| Cell-Cell communication |
| Interleukin-4 and Interleukin-13 signaling |
| G alpha (i) signalling events |
| Translocation of SLC2A4 (GLUT4) to the plasma membrane |
| MAP kinase activation |
| Signaling by GPCR |
| TNFR2 non-canonical NF-kB pathway |
| RHOG GTPase cycle |
| ESR-mediated signaling |

| Pathway |
| --- |
| RAC1 GTPase cycle |

Table 5: Pathways uniquely identified by PseudotimeDE

| Pathway |
| --- |
| HIV Infection |
| mRNA Splicing |
| mRNA Splicing - Major Pathway |
| HIV Life Cycle |

Table 6: Common pathways identified by both methods

| Pathway |
| --- |
| Eukaryotic Translation Elongation |
| Peptide chain elongation |
| Formation of a pool of free 40S subunits |
| GTP hydrolysis and joining of the 60S ribosomal subunit |
| Viral mRNA Translation |
| Selenocysteine synthesis |
| L13a-mediated translational silencing of Ceruloplasmin expression |
| Eukaryotic Translation Termination |
| Nonsense Mediated Decay (NMD) independent of the Exon Junction Complex (EJC) |
| Eukaryotic Translation Initiation |
| Cap-dependent Translation Initiation |
| SRP-dependent cotranslational protein targeting to membrane |
| Selenoamino acid metabolism |
| Response of EIF2AK4 (GCN2) to amino acid deficiency |
| Influenza Viral RNA Transcription and Replication |
| Influenza Infection |
| Metabolism of amino acids and derivatives |
| Major pathway of rRNA processing in the nucleolus and cytosol |
| Translation |
| rRNA processing |
| rRNA processing in the nucleus and cytosol |
| Infectious disease |
| Viral Infection Pathways |
| Disease |
| Metabolism of RNA |

#### 3.4 EnrichR Knowledge Network

We constructed knowledge graph (KG) networks using Enrichr's Mammalian Phenotype 2017 enrichment Smith, Goldsmith, and Eppig (2004) showed in figure 9. Despite relying on a smaller subset of differentially expressed genes, scFPC-DE recovered a comparably rich phenotypic profile (top panel), capturing B cell–specific phenotypes such as abnormal B cell differentiation, altered physiology, and immunoglobulin level abnormalities. In contrast, PseudotimeDE (bottom panel) identified a larger gene set, resulting in a similar network. These results shows scFPC-DE's efficiency in obtaining functionally meaningful gene–phenotype associations with fewer inputs and complement the results discussed in the main text (Figure 7d–e).

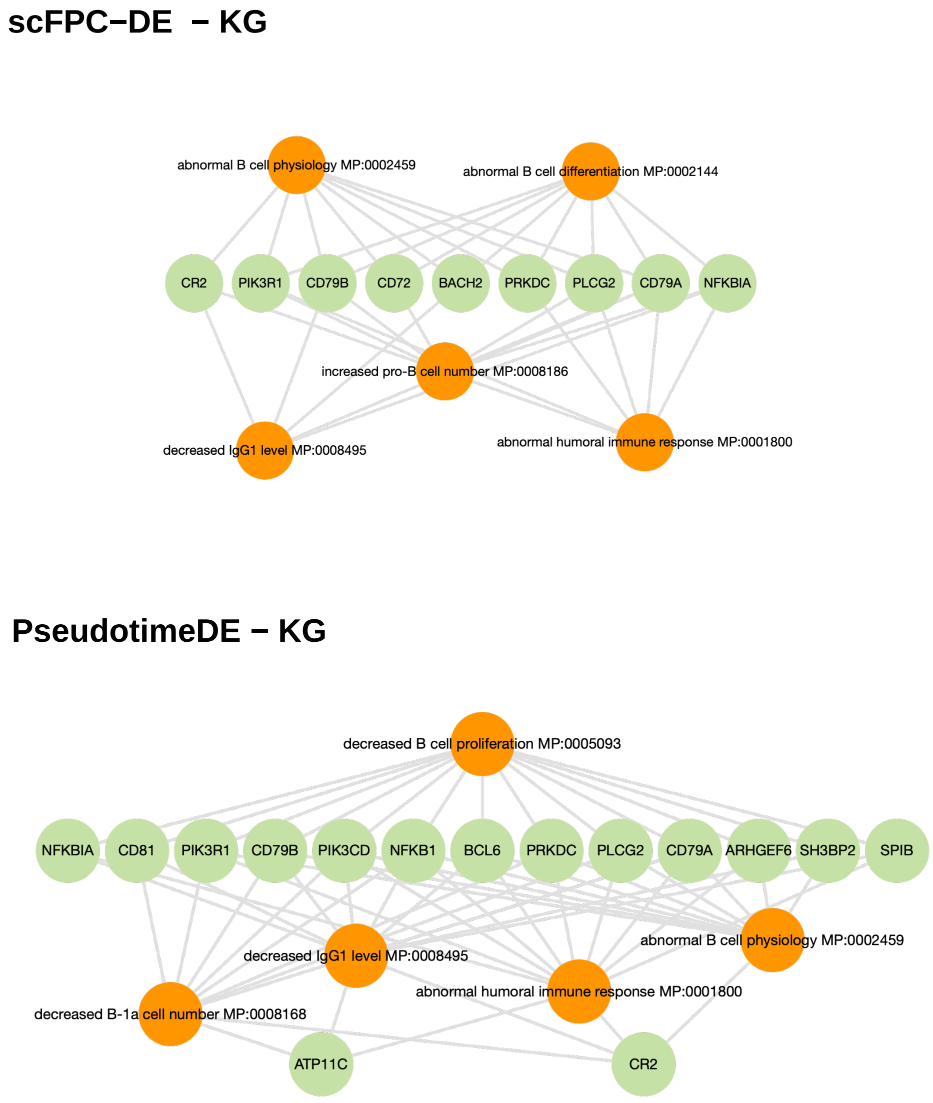

Figure 9: Knowledge graph (KG) networks from Enrichr illustrate phenotypic enrichment based on gene sets identified by scFPC-DE (top) and PseudotimeDE (bottom), using the MGI Mammalian Phenotype Level 4 2021 ontology. Despite relying on a more compact set of differentially expressed genes, scFPC-DE recovers key immune-related pathways that also appear in the broader PseudotimeDE results.

Ramsay, James O, and Bernard W Silverman. 2005. *Functional Data Analysis*. Book. Springer.

Ramsay, James, Giles Hooker, Spencer Graves, JO Ramsay, Giles Hooker, and Spencer Graves. 2009. "Introduction to Functional Data Analysis." Journal Article. *Functional Data Analysis with R and MATLAB*, 1–19.

Shang, Han Lin. 2014. "A Survey of Functional Principal Component Analysis." Journal Article. *AStA Advances in Statistical Analysis* 98: 121–42.

Smith, Cynthia L, Carroll-Ann W Goldsmith, and Janan T Eppig. 2004. "The Mammalian Phenotype Ontology as a Tool for Annotating, Analyzing and Comparing Phenotypic Information." Journal Article. *Genome Biology* 6 (1): R7.

Song, Dongyuan, and Jingyi Jessica Li. 2021. "PseudotimeDE: Inference of Differential Gene Expression Along Cell Pseudotime with Well-Calibrated p-Values from Single-Cell RNA Sequencing Data." Journal Article. *Genome Biology* 22 (1): 124.

Song, Dongyuan, Qingyang Wang, Guanao Yan, Tianyang Liu, Tianyi Sun, and Jingyi Jessica Li. 2024. "scDesign3 Generates Realistic in Silico Data for Multimodal Single-Cell and Spatial Omics." Journal Article. *Nature Biotechnology* 42 (2): 247–52.

Stewart, Alexander, Joseph Chi-Fung Ng, Gillian Wallis, Vasiliki Tsioligka, Franca Fraternali, and Deborah K Dunn-Walters. 2021. "Single-Cell Transcriptomic Analyses Define Distinct Peripheral b Cell Subsets and Discrete Development Pathways." Journal Article. *Frontiers in Immunology* 12: 602539.

Yu, Guangchuang, and Qing-Yu He. 2016. "ReactomePA: An r/Bioconductor Package for Reactome Pathway Analysis and Visualization." Journal Article. *Molecular BioSystems* 12 (2): 477–79.
